# Multiscale oocyst wall mechanics govern coccidian resistance

**DOI:** 10.64898/2026.06.23.734144

**Authors:** Jana El Husseiny, Rémy Torro, Laura Sedano, Catherine Cazeaux, Hugo Le Guenno, Artemis Kosta, Mickaël Riou, Jean-Michel Répérant, Stéphanie La Carbona, Anne Silvestre, Julien Husson, Pierre-Henri Puech, Aurélien Dumètre

**Affiliations:** Aix Marseille University, CNRS, INSERM, LAI, Marseille, France; Hydrodynamics laboratory (LadHyX), CNRS, École polytechnique, Institut Polytechnique de Paris, 91120 Palaiseau, France; INRAE, UMR ISP, Université de Tours, Nouzilly 37380, France; ACTALIA, 310 Rue Popielujko, Saint-Lô, 50000, France; Aix Marseille University, CNRS, IMM, Microscopy Core Facility, IM2B, Marseille, France; INRAE, UE-1277 Plateforme d’Infectiologie expérimentale (PFIE), Centre Val de Loire, 37380 Nouzilly, France; Avian and Rabbit Virology, Immunology, Parasitology Unit, ANSES, Laboratory of Ploufragan-Plouzané-Niort, 22440 Ploufragan, France

## Abstract

Coccidian parasites spread through oocysts, environmental transmission stages that must survive harsh conditions yet rupture on cue inside the host to release infectious sporozoites. How the oocyst wall reconciles these opposing demands is unknown. Using single-oocyst microindentation, quantitative imaging and infection assays in *Eimeria acervulina* and *Eimeria tenella*, we show that the bilayered wall is functionally partitioned: the chemically fragile outer layer is mechanically dispensable, while the inner layer bears the mechanical load and remains impermeable to macromolecules and limiting osmotic exchange even after the outer layer is stripped. Oocysts behave as pressurised shells that fail preferentially at the anterior pole, a fixed mechanical weak point that channels sporocyst release regardless of where force is applied. Wall structure, autofluorescence, mechanics and infectivity prove dissociable: bleach removes the outer layer without weakening the wall or fully abolishing infectivity, whereas heat weakens the wall and abolishes infectivity while leaving the bilayered structure intact. These findings define a mechanical logic for coccidian oocyst resistance, disinfection and controlled excystation, and identify the inner wall as the critical target for inactivating environmentally transmitted parasites.

## Introduction

Coccidia are a large group of intracellular parasites (∼2,000 species) transmitted worldwide through oocysts, contaminating soil, water and food ^1^. Environmental oocysts contain infective sporozoites enclosed within sporocyst and oocyst walls ^2^. Upon ingestion, sporozoites are released through excystation and invade the intestinal epithelium, where asexual replication and sexual reproduction produce oocysts that are shed in faeces. Outside the host, these oocysts undergo sporulation within hours to days, depending on species and conditions, to form infective sporozoites. Most coccidia are host-specific, including the veterinary-relevant chicken *Eimeria* species ^3^ and the human pathogen *Cyclospora cayetanensis* ^4^, and remain confined to the digestive tract where they can cause diarrhoeal disease. A notable exception is *Toxoplasma gondii*, whose oocysts are produced only by felids but which can infect a wide range of birds and mammals, including humans. In immunocompetent hosts, infection is usually asymptomatic or causes only mild, self-limiting illness; severe cerebral and ocular disease arises mainly in immunocompromised individuals such as HIV patients and the developing foetus ^5^. In the absence of specific drugs and affordable vaccines in many settings ^6,7^, understanding how oocysts resist environmental conditions while retaining infectivity is a key step towards limiting the public health and economic burden of coccidian infections.

*In vitro* and *in vivo* studies showed that coccidian oocysts remain infectious across a broad range of environmental conditions, including wide changes in temperature and osmolarity ^8^. They also remain infective upon treatment with common detergents and chemical disinfectants, limiting standard decontamination strategies ^9^. Complete inactivation typically requires prolonged desiccation, lasting from days to weeks depending on relative humidity, or sustained freezing at −21 °C for at least one day, or heating exceeding 60 °C for several minutes, conditions that are difficult to implement for water and food decontamination in domestic and industrial settings ^8,10,11^. These observations point to the oocyst wall and secondarily to the sporocyst wall as key determinants of sporozoite infectivity. These walls must remain tightly sealed and impermeable in the external environment to protect the parasites, yet be efficiently opened within the host digestive tract at the right moment and place to release the sporozoites and therefore initiate infection ^3,12^. This dual requirement for environmental resistance and timely opening is fundamental to coccidian biology and directly relevant to infection control in humans and livestock, yet the mechanisms governing wall rupture remain poorly understood.

A general structural and molecular model proposes that both walls are bilayered and protein-rich (>90%), with triglycerides covering their surface ^13–15^. The core structure of the walls is dominated by tyrosine-rich proteins forming dityrosine and DOPA–protein cross-links, which confer a characteristic blue autofluorescence under ultraviolet excitation ^16^, a feature that can be used as a quantitative proxy for structural changes in the wall ^17^. The inner oocyst wall layer is reinforced by a network of β-1,3-glucan fibrils, proposed to provide structural rigidity ^18^. However, this architecture does not account for the functional contribution of each layer to wall integrity under chemical and physical stress, nor for the detailed mechanisms of their opening in response to host digestive cues ^3,12^.

A key finding of previous studies is that treatment of *T. gondii* oocysts with household bleach selectively removes the outer wall layer (∼20 nm thick) without affecting the inner wall (∼70 nm thick) or the sporocyst wall, while preserving sporozoite infectivity ^17^. Elasticity measurements using atomic force microscopy (AFM) showed that the inner wall layer exposed at the oocyst surface following bleach treatment is mechanically as robust as the intact bilayer wall, with a measured Young’s modulus of 1–10 MPa, while exhibiting a larger adhesion capacity ^17^. These findings suggest that the two oocyst wall layers may be functionally distinct and that wall mechanics, measured to be as stiff as soft elastomers or plant cell walls ^19,20^, may be critical for protecting oocyst content from external stressors while regulating the release of sporozoites in the host digestive tract.

In this study, we used *Eimeria acervulina* and *Eimeria tenella*, two chicken coccidia amenable to experimental infection and large-scale oocyst production, as surrogate models to investigate coccidian oocyst wall mechanobiology ^21^. The two species infect distinct regions of the chicken gut, with the moderately pathogenic *E. acervuline* colonising the proximal small intestine, and the more pathogenic *E. tenella*, the caeca, implying either prolonged intestinal transit or bloodstream-mediated transport of sporozoites to the target site ^22,23^. To investigate oocyst wall mechanobiology, we employed a multi-modal approach combining microindentation, atomic force microscopy (AFM), quantitative autofluorescence imaging with deep learning-based segmentation, transmission electron microscopy (TEM), lectin-binding and osmotic permeability assays and *in vivo* infectivity assessment. We demonstrate that the inner wall layer confers the mechanics and impermeability of the oocyst wall, identify the anterior pole of oocysts as an intrinsically, mechanically weak site that channels sporocyst release and establish that chemical oxidising treatments erode the outer layer without completely eliminating the infectivity of the sporozoites. Together, these findings establish a functional architecture for *Eimeria* oocysts, provide a conceptual framework for understanding oocyst wall function across coccidia, and identify the wall-opening step preceding infection as a potential intervention point.

## Results

### Digestive factors leave oocyst morphology and wall structure largely intact

Trypsin and bile salts are key components of *in vitro* excystation protocols, routinely used to simulate the digestive environment encountered by coccidian oocysts within the host gut ^12^. Whether these factors directly remodel the oocyst wall and act as triggers for wall opening and sporozoite release remains unclear. To address this, we exposed *E. acervulina* and *E. tenella* oocysts to trypsin, bile salts, or their combination, and quantified changes in oocyst morphology, wall autofluorescence (AF), and ultrastructure.

*E. acervulina* oocysts retained their ovoid morphology after all treatments, with no visible wall disruption or ruptured oocysts in the parasite suspensions (Fig. 1a). Trypsin alone did not alter oocyst area, which we define as the cross-sectional area observed in microscopy, while bile salt-containing treatments produced a small increase (Fig. 1b; Cliff’s δ are reported in Supplementary Table 1). Wall AF declined progressively with digestive challenge, from no change with bile salts to a small reduction with trypsin and a large reduction with the combined treatment (Fig. 1c and Supplementary Table 1). *E. tenella* oocysts similarly preserved their overall apparent structure and shape (Fig. 1d), and neither trypsin, bile salts, nor their combination altered oocyst area (Fig. 1e). Wall AF was broadly comparable between conditions, although the combined trypsin and bile salt treatment produced a small increase compared to the untreated control (Fig. 1f and Supplementary Table 1).

**Figure 1.**
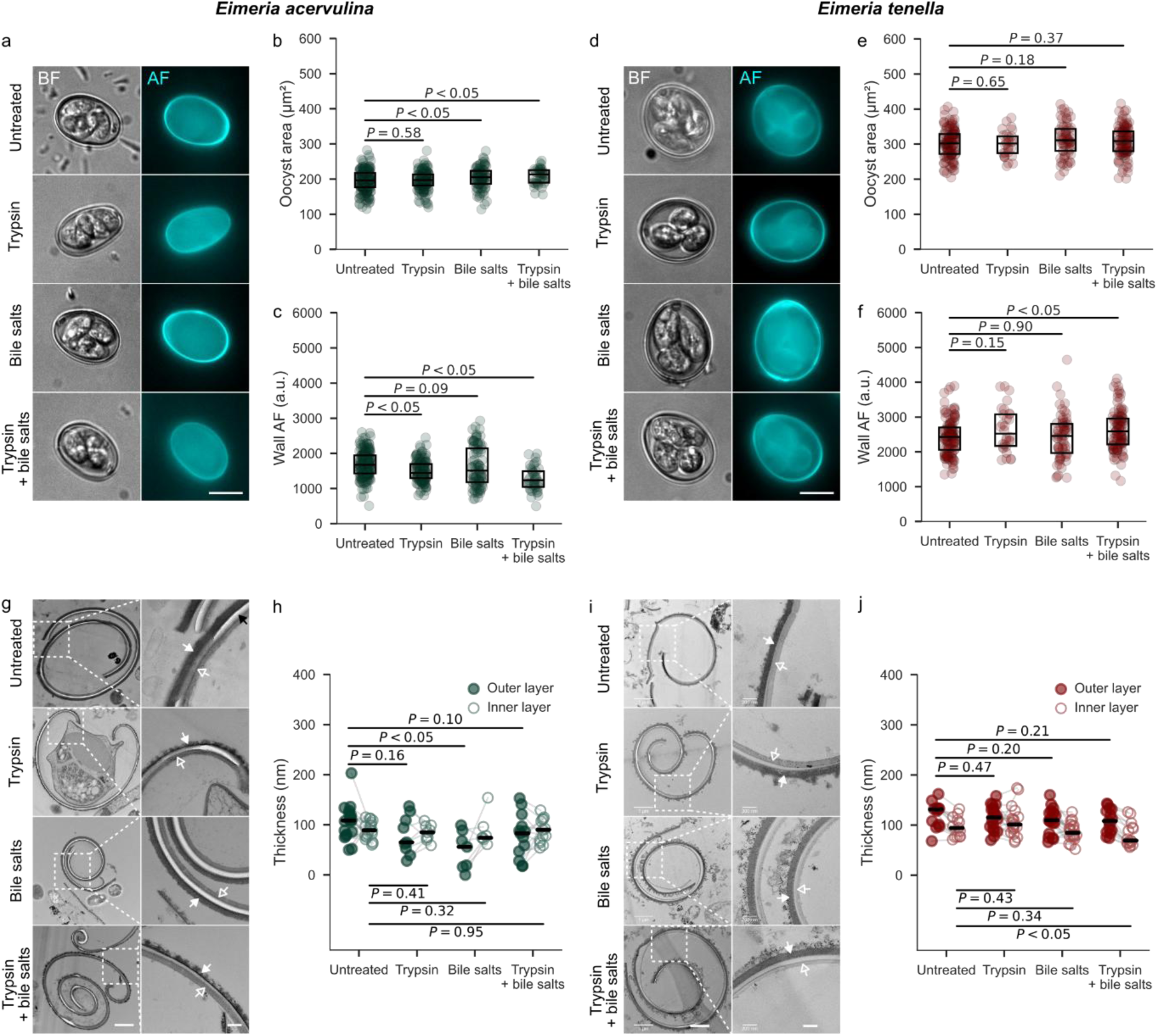
| Oocyst morphology and wall structure are largely preserved after exposure to digestive factors. **a,** Representative bright-field (BF) and autofluorescence (AF) images of *E. acervulina* oocysts under untreated, trypsin, bile salts, and trypsin + bile salts conditions. Scale bars, 10 µm. **b,** Quantification of oocyst area, cross-sectional area measured from microscopy images, in *E. acervulina* across treatment conditions. **c,** Quantification of oocyst wall AF intensity in *E. acervulina*. a.u., arbitrary units. Each dot represents an individual oocyst; boxes indicate the interquartile range and horizontal lines indicate the median. **d–f,** Same as **a–c** for *E. tenella* oocysts. **g,** Representative transmission electron micrographs of *E. acervulina* oocyst wall ultrastructure under all four conditions. Scale bars, 1 µm in overview images and 200 nm in magnified views. White arrows indicate the outer wall layer, open arrows indicate the inner wall layer, and black arrows indicate imaging artefacts. **h,** Outer- and inner-wall thickness in *E. acervulina*. Filled and open circles indicate the outer and inner layers, respectively. Each dot represents an individual measurement and horizontal bars indicate the median. **i,j,** Same as **g,h** for *E. tenella* oocysts. *P* values were calculated using two-tailed Mann–Whitney *U* tests.

TEM revealed a characteristic bilayered ultrastructure of the coccidian oocyst wall, with each layer approximately 100 nm thick, separated by an electron-lucent interspace of variable width likely attributable to sample preparation artefacts, giving an estimated total wall thickness of ∼200 nm. Neither species showed disruption of this bilayered organisation following any treatment (Fig. 1g,i). Quantification of wall-layer thickness showed that in *E. acervulina*, digestive factors produced modest alterations overall, although bile salt treatment reduced outer-layer thickness by approximately 50%, while the inner layer remained unaffected (Fig. 1h). In *E. tenella*, the combined trypsin and bile salt treatment modestly reduced inner-layer thickness, while the outer layer and individual treatments produced no significant changes (Fig. 1j). Together, these results show that trypsin and bile salts do not visibly rupture intact *E. acervulina* and *E. tenella* oocysts and largely preserve wall bilayer organisation, while inducing modest, species-and layer-specific reductions in wall thickness.

### Chemical disinfectants selectively erode the outer oocyst wall layer but fail to completely abolish parasite infectivity

Having established that digestive factors at physiological concentrations have little impact on the oocyst wall structure, we next asked whether harsher chemical and physical treatments, including bleach, ozone, and heat, produce more pronounced structural changes as observed for *T. gondii* ^17^.

*E. acervulina* oocysts retained their ovoid shape and did not open after all treatments, although AF signal was visibly reduced after bleach and ozone exposure (Fig. 2a). Bleach and heat reduced oocyst area, whereas ozone had negligible effect (Fig. 2b and Supplementary Table 1). Wall AF was decreased by all three treatments, with the strongest reduction recorded after bleach and ozone (Fig. 2c). Similarly, in *E. tenella*, oocysts remained morphologically intact following all treatments (Fig. 2d). Oocyst area was reduced by heat and ozone, whereas bleach had essentially negligible effect (Fig. 2e and Supplementary Table 1). Strikingly, heat treatment increased wall AF, while bleach and ozone reduced the signal (Fig. 2f).

**Figure 2.**
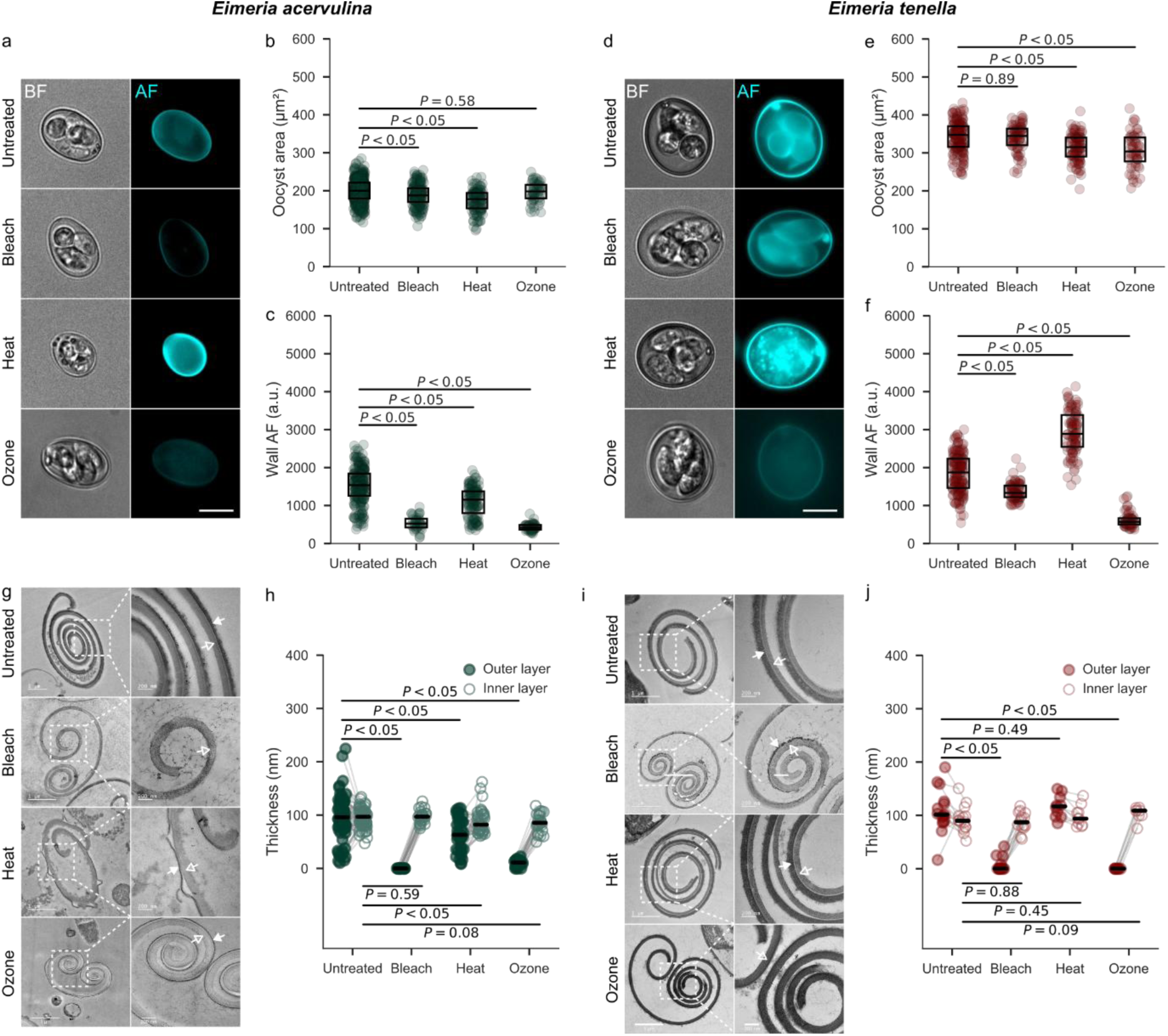
| Harsh chemical and thermal treatments differentially remodel the wall architecture of *Eimeria* oocysts. **a,** Representative bright-field (BF) and autofluorescence (AF) images of *E. acervulina* oocysts under untreated, bleach, heat, and ozone conditions. Scale bars, 10 µm. **b,** Quantification of oocyst area in *E. acervulina* across treatment conditions. **c,** Quantification of oocyst wall AF intensity in *E. acervulina*. a.u., arbitrary units. Each dot represents an individual oocyst; boxes indicate the interquartile range and horizontal lines indicate the median. **d–f,** Same as **a–c** for *E. tenella*. **g,** Representative transmission electron micrographs of *E. acervulina* oocyst wall ultrastructure under all four conditions. Scale bars, 1 µm in overview images and 200 nm in magnified views. White arrows indicate the outer wall layer and open arrows indicate the inner wall layer. **h,** Outer- and inner-wall thickness in *E. acervulina*. Filled and open circles indicate the outer and inner layers, respectively. Each dot represents a measurement from a single oocyst and horizontal bars indicate the median. **i,j,** Same as **g,h** for *E. tenella*. *P* values were calculated using two-tailed Mann–Whitney *U* tests.

TEM images revealed treatment-dependent modifications of the oocyst wall ultrastructure, with the most prominent changes affecting the outer wall layer (Fig. 2g,i). In *E. acervulina*, bleach caused complete loss of the outer layer, whereas ozone produced variable outer-layer erosion, ranging from near-complete removal to retention of a markedly thinned layer; quantification confirmed that all three treatments significantly reduced outer-layer thickness, with heat also reducing inner-layer thickness (Fig. 2h). In *E. tenella*, bleach and ozone similarly induced visible outer-layer loss and significantly reduced outer-layer thickness, whereas heat did not measurably affect the outer layer and none of the treatments altered inner-layer thickness (Fig. 2j). Despite these outer-layer alterations, the inner wall layer was not removed and retained a homogeneous appearance across treatments, indicating that bleach and ozone preferentially compromise the outer oocyst wall without causing complete wall collapse.

We next investigated whether the observed changes in the wall structure translated to loss of oocyst infectivity. For this, we orally inoculated chickens with untreated, bleach-, ozone- or heat-treated oocysts from *E. tenella*, the more pathogenic of the two species studied ^6^. Oocyst infectivity was quantified by scoring caecal lesions associated with parasite development and by enumerating newly formed oocysts in the caecal contents of chickens euthanised 7 days post-inoculation (Supplementary Fig. 1a). As expected, heat-treated oocysts produced no lesions and no oocyst output in any animal (Supplementary Fig. 1b,c). Ozone treatment significantly reduced both lesion severity and oocyst output compared with untreated controls, whereas bleach produced no significant reduction in oocyst output (Supplementary Fig. 1b,c).

Together, these results show that bleach, ozone, and heat differentially remodel oocyst wall architecture while largely preserving overall oocyst morphology. However, their effects on infectivity diverge markedly, indicating that ultrastructural alterations of the wall are not necessarily predictive of parasite infectivity.

### Outer-wall disruption exposes glycan motifs without breaching the inner layer

Electron microscopy established that the outer and inner wall layers respond differentially to external factors, with the outer layer being selectively eroded by bleach and ozone while the inner layer remains structurally intact across all treatments. However, whether this structural hierarchy translates into distinct surface chemistry and wall porosity could not be resolved by ultrastructural imaging alone. To address this, we quantified lectin accessibility to the different wall substructures using TRITC-conjugated lectins on untreated, bleach-, heat-, and ozone-treated *E. acervulina* and *E. tenella* oocysts.

WGA (∼36 kDa) and MPL (∼40 kDa) are relatively small lectins that recognise GlcNAc-and Gal/GalNAc-containing structures, respectively. WGA-reactive targets have been reported on the oocyst wall surface in *E. acervulina* and *T. gondii* ^24^. MPL-reactive glycoproteins have been localised to both the inner oocyst wall and the sporocyst wall of *E. tenella* and *T. gondii* ^18,25^. PNA (∼110 kDa), a larger tetrameric lectin that recognises Galβ1–3GalNAc motifs, was included because PNA-reactive glycoconjugates have been reported in *E. tenella* oocysts and sporozoites, although their precise accessibility within intact oocyst wall layers is less well defined ^26,27^. Thus, PNA was used here to test whether Galβ1–3GalNAc-containing targets are accessible from the exterior of the oocyst wall or restricted to the inner wall face.

In untreated whole oocysts, WGA, MPL, and PNA labelling was weak in both species (Fig. 3a,d). Heat-treated oocysts showed equally limited labelling across all three lectins (Fig. 3a–f). Bleach treatment, which selectively removes the outer wall layer, produced a marked increase in peripheral WGA and MPL fluorescence in both species (Fig. 3a–f), reaching approximately 327-fold and 596-fold above unstained controls for MPL in *E. acervulina* (Fig. 3c) and *E. tenella* (Fig. 3f), respectively. Unexpectedly, ozone produced a distinct pattern: MPL signals remained comparable to those of untreated oocysts, while WGA signals were reduced below untreated levels in both *Eimeria* species (Fig. 3a–f). In all labelled oocysts across treatments, fluorescence was restricted to the oocyst periphery, with no detectable staining of internal sporocysts or sporozoites.

**Figure 3.**
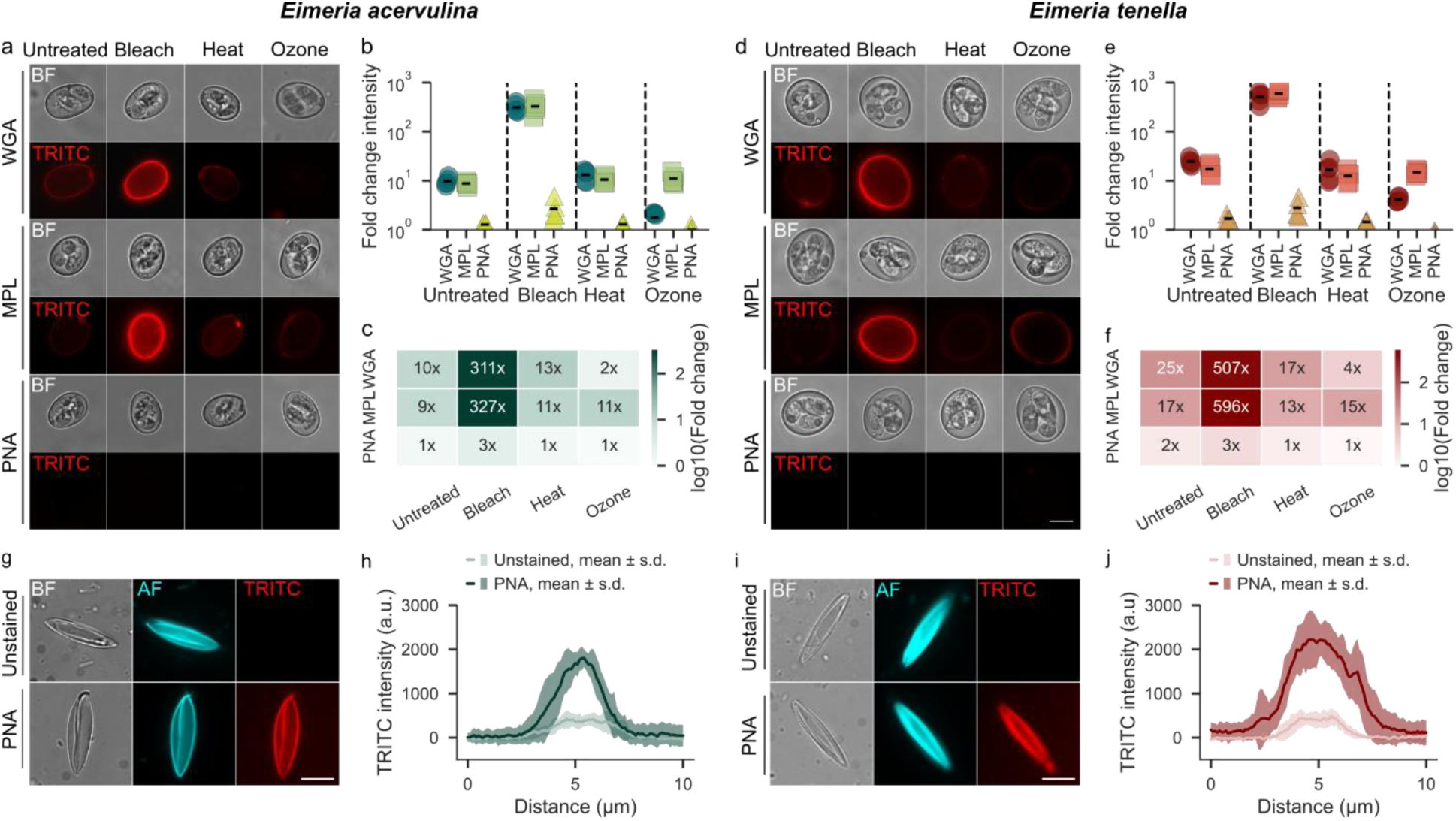
| Outer-wall disruption exposes inner-wall lectin-binding sites without permitting access to the oocyst interior in *E. acervulina* and *E. tenella*. **a**, Representative bright-field (BF) and TRITC fluorescence images of *E. acervulina* oocysts stained with wheat germ agglutinin (WGA), *Maclura pomifera* lectin (MPL), or peanut agglutinin (PNA) after bleach, heat, or ozone treatment, alongside untreated controls. Scale bars, 10 µm. **b,** Quantification of lectin-associated fluorescence intensity from flow cytometry, expressed as fold change relative to unstained controls, for WGA, MPL, and PNA in *E. acervulina*. **c,** Heat map showing lectin-associated fluorescence fold change in *E. acervulina*. Numeric values indicate fold change relative to unstained controls; colour intensity represents log₁₀-transformed fold change. **d–f,** Same as **a–c** for *E. tenella*. **g,** Representative BF, autofluorescence (AF), and TRITC images of sonicated *E. acervulina* oocysts under unstained and PNA-stained conditions. Scale bars, 10 µm. **h,** Mean TRITC fluorescence intensity profiles measured across the transverse axis of sonicated *E. acervulina* oocysts under unstained and PNA-stained conditions. Solid lines indicate the mean and shaded areas indicate s.d. **i,j,** Same as **g,h** for *E. tenella*. Data points represent independent experimental replicates; horizontal bars indicate the median. Line profile plots were obtained from seven oocysts per condition.

To test whether PNA-reactive motifs were present within the oocyst wall, oocysts were mechanically disrupted by sonication, an efficient method for breaching the intact wall and exposing internal surfaces that are otherwise inaccessible from the exterior. Sonicated oocysts displayed a clear PNA signal, with fluorescence intensity profiles revealing enrichment at the inner face of the wall (Fig. 3g–j). These data demonstrate that PNA-reactive molecules are confined to the inner wall face and are inaccessible from the exterior regardless of treatment.

To further investigate oocyst wall permeability to molecules smaller than lectins, including ions and water, we next quantified wall structural integrity or dimensions under various osmotic conditions. These conditions were chosen to span the range of osmotic challenges that coccidian oocysts may encounter in the environment, including sandy soils, saline waters and freshwater. Oocysts were exposed for 24 h to near-isosmotic 1× PBS, a standard cellular buffer, hyperosmotic 10× PBS, hypoosmotic water, or recovery conditions after hyperosmotic exposure (10× PBS → 1× PBS). Surface buckling was observed in some *E. acervulina* oocysts, whereas *E. tenella* oocysts showed no apparent morphological changes (Supplementary Fig. 2a,d). Oocyst area remained unchanged in both species under all conditions (Supplementary Fig. 2b,e), indicating that the oocyst wall neither collapsed nor swelled passively under osmotic gradients. Osmotic challenge reproduced this species divergence in wall AF: AF decreased in *E. acervulina* across all conditions but was unchanged or increased in *E. tenella* (Supplementary Fig. 2c,f and Supplementary Table 1), recapitulating the species-specific responses observed after chemical treatment.

Together, these results show that the inner-wall glycan motifs are unmasked following bleach-mediated removal of the outer layer, whereas ozone induces a chemically distinct wall state in which these lectin-reactive molecules remain poorly detectable, despite comparable outer-layer erosion. PNA exclusion under all treatments and the absence of osmotic volume change establish the inner wall as a chemically selective, macromolecule-tight barrier whose impermeability is preserved after outer-layer loss.

### Oocyst wall mechanics reveals orientation-dependent rupture and treatment-specific weakening

To investigate wall mechanics, we directly quantified wall rupture mechanics at the single-oocyst level using microindentation, probing how indentation orientation, digestive factors, and disinfection treatments each modulate the forces required to breach the wall.

Atomic force microscopy (AFM) has previously been applied to *T. gondii* oocysts to evaluate wall mechanical properties, although a similar approach based on immobilising *Eimeria* oocysts on poly-L-lysine-coated substrates proved inconsistent, precluding systematic measurements (Supplementary Fig. 3a–c). AFM measurements on a handful of successfully immobilised oocysts nevertheless revealed treatment-dependent differences in surface adhesion and apparent local wall stiffness: bleach- and heat-treated oocysts displayed increased adhesion forces and decreased local Young’s moduli of the oocyst wall compared with untreated controls (Supplementary Fig. 3d,e). Young’s moduli of the wall were in the MPa range, and bleach rendered the oocyst more adhesive, consistent with observations reported for *T. gondii* ^17^. Aside with *T. gondii*, we were unable to pierce the oocyst wall to analyse its resistance to force using our AFM setup.

We therefore developed a micropipette-based microindentation assay to measure wall rupture forces on individual oocysts without having to adhere them. Oocysts were held away from the surface by gentle aspiration with a stiff micropipette and indented with needle-shaped calibrated flexible micropipettes until wall failure, evidenced by a sharp drop in the force–time curve consistent with the observations made simultaneously in bright field (Fig. 4a,b and Supplementary Video 1) ^28^. Rupture force was defined as the maximum force recorded at the point of failure, measured relative to the baseline force with the indenter far from the oocyst (Fig. 4a), and reflects the combined contributions of wall material properties and, where present, internal pressure.

**Figure 4.**
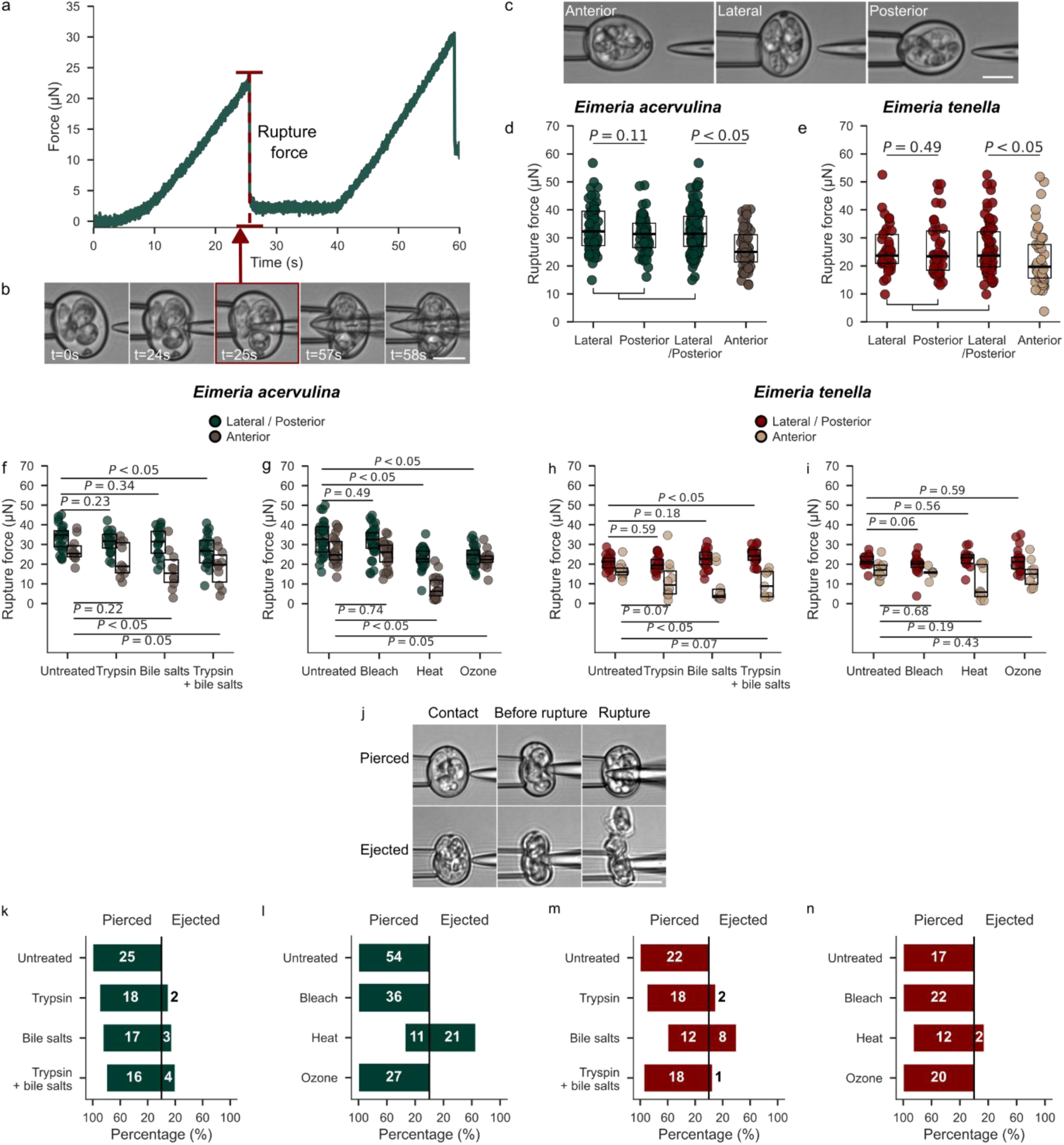
| Microindentation reveals orientation-dependent rupture forces and treatment-specific wall failure in *Eimeria* oocysts. **a**, Representative force–time curve obtained during microindentation of a single oocyst. Force increases until rupture, after which a sharp force drop indicates mechanical failure of the oocyst wall. The rupture force is indicated by the red dashed line and arrow. The second peak corresponds to the distal wall rupture. **b,** Bright-field time-lapse images of an *E. acervulina* oocyst during microindentation at the indicated time points. The red box and arrow highlight the frame corresponding to the rupture of the proximal wall. Scale bar, 10 µm. **c,** Representative bright-field images illustrating the three indentation orientations used: anterior, lateral and posterior. Scale bar, 10 µm. **d,e,** Rupture force of untreated *E. acervulina* (**d**) and *E. tenella* (**e**) oocysts indented at the lateral, posterior, pooled lateral/posterior or anterior positions. **f–i,** Rupture force of *E. acervulina* (**f,g**) and *E. tenella* (**h,i**) oocysts after exposure to digestive factors (**f,h**) or disinfectant treatments (**g,i**), compared with untreated controls. Oocysts were indented at the lateral/posterior position or at the anterior pole, as indicated by the colour code. In **d–i**, boxes indicate the interquartile range and horizontal lines indicate the median; each dot represents an individual oocyst. *P* values were calculated using two-tailed Mann–Whitney *U* tests. **j,** Representative bright-field time-lapse images illustrating the two modes of wall failure observed during microindentation, both with the probe applied at the lateral position. Three successive frames are shown for each: contact, before rupture, and rupture. Top: direct piercing, in which wall failure occurs at the lateral contact point. Bottom: anterior ejection, in which wall failure occurs preferentially at the anterior pole, resulting in sporocyst expulsion through the anterior pole despite force being applied laterally. Scale bar, 10 µm. **k–n,** Percentage of oocysts that were pierced or showed content ejection during microindentation after exposure to digestive factors in *E. acervulina* (**k**) and *E. tenella* (**m**), or after disinfectant treatments in *E. acervulina* (**l**) and *E. tenella* (**n**). Numbers inside bars indicate the number of oocysts in each category.

We first asked whether rupture force depended on indentation orientation due to the anisotropy of the oocyst. Indenting untreated oocysts at the anterior pole (the more pointed, tapered end of the oocyst), posterior pole, or lateral side (Fig. 4c and Supplementary Video 2) revealed that the anterior pole consistently required lower forces to rupture than lateral or posterior positions, which behaved similarly, in both species (Fig. 4d,e). As lateral and posterior rupture forces were statistically indistinguishable, these positions were pooled for all subsequent analyses. Median rupture forces were approximately 25 µN anteriorly versus 32 µN laterally/posteriorly in *E. acervulina*, and 20 µN versus 24 µN in *E. tenella*, indicating that the oocyst wall is mechanically anisotropic.

We next tested whether digestive factors altered wall mechanical resistance. Trypsin alone had no effect on rupture force in either species, whereas bile salts reduced rupture force at the anterior pole in both species (large effect, Fig. 4f,h and Supplementary Table 1). The combined trypsin–bile salt treatment had opposite effects at the lateral/posterior positions: in *E. acervulina* it modestly reduced rupture force, whereas in *E. tenella* it increased it (Fig. 4f,h and Supplementary Table 1).

Disinfection treatments produced stronger mechanical changes than digestive factors, with species-specific outcomes. In *E. acervulina*, heat and ozone markedly reduced rupture force at both lateral/posterior and anterior positions, whereas bleach-treated oocysts showed rupture forces comparable to untreated controls (Fig. 4g and Supplementary Table 1), consistent with bleach acting selectively on the outer wall layer while the inner layer remains intact. In *E. tenella*, none of the disinfection treatments produced statistically significant changes in rupture force at any indentation position (Fig. 4i), indicating that the wall of this species appears to be mechanically more resilient to the treatments tested.

Microindentation also revealed two distinct modes of wall failure. In most oocysts, indentation produced complete wall rupture at the contact site. However, in a subset of oocysts, particularly after heat treatment, force applied at lateral or posterior positions caused localised failure specifically at the anterior pole, with sporocysts expelled through this site rather than at the point of contact (Fig. 4j and Supplementary Video 3). This remote failure mode directly demonstrates that the anterior pole represents a mechanically weaker region of the wall. The frequency of sporocyst ejection versus complete rupture varied by species and treatment condition (Fig. 4k-n), with heat-treated *E. acervulina* showing the highest ejection frequency. This is consistent with the reduced rupture forces recorded at this site after heat treatment (Fig. 4g).

The lectin permeability assays established that the inner wall constitutes a highly selective barrier, inaccessible to macromolecules from the exterior, even after outer-layer removal. A corollary of this impermeability would be that the internal pressure of the oocyst sealed within the intact inner wall cannot be modulated by changes in external osmolarity. To test whether external osmolarity modulates rupture force, we measured wall mechanics after osmotic treatments. Consistent with the impermeability of the inner wall, rupture force was unaffected in *E. acervulina* and, in *E. tenella*, was only moderately increased at lateral/posterior positions across all conditions; no sporocyst ejection was observed in either species (Supplementary Fig. 2g–j). These results confirm that external osmolarity does not perturb internal pressure within the range and over the timescale of the tested treatments, consistent with an inner layer whose integrity is maintained independently of external ionic conditions.

To quantify the internal pressure of the oocyst, we estimated it from the relationship between contact stiffness and local oocyst radius using a pressure-dominated shell model (*k* = *πPR_eq_,* where *P* is the internal pressure and *R_eq_* an equivalent radius – an average local radius of curvature; Supplementary Fig. 4a) ^29^. Contact stiffness scaled linearly with equivalent radius across indentation orientations, consistent with pressurized-shell mechanics (Supplementary Fig. 4b-e). Under untreated conditions, estimated internal pressure was approximately 124 kPa in *E. acervulina* and 71 kPa in *E. tenella*. Bleach left estimated internal pressure essentially unchanged in *E. acervulina* but reduced it modestly in *E. tenella*, whereas heat and ozone decreased the estimated pressure. Osmotic perturbations had only a small effect on estimated internal pressure: the most extreme hyperosmotic challenge (10× PBS) reduced it by ∼15–20% in both species, whereas hypo-osmotic water left it unchanged (Supplementary Fig. 4f,g).

Together, these results show that oocyst rupture mechanics are orientation-dependent, with the anterior pole acting as the mechanically weakest region, and that wall resistance is modulated by treatment in a species-specific manner. They further indicate that the oocyst behaves as a pressurised, mechanically anisotropic shell whose rupture properties are largely maintained after outer-layer loss but weakened by heat or ozone in *E. acervulina*.

## Discussion

This study establishes a functional mechanical framework for the coccidian oocyst wall, revealing a hierarchy of layer-specific properties that enables these structures to withstand environmental stress while preserving the capacity for efficient excystation. By combining structural imaging and infectivity assays with direct force measurements up to wall failure, we identify three key mechanistic determinants. First, the inner wall layer governs the mechanical resistance and permeability of the oocyst wall. Second, this wall contains a spatially fixed weak point at the anterior pole that directs sporocyst release under localised force, independent of where that force is applied. Third, structural or chemical alterations of the wall do not predict complete loss of sporozoite infectivity.

The first determinant is that the inner wall layer carries the mechanical and permeability functions of the oocyst wall. Strikingly, complete removal of the outer wall layer by bleach left rupture forces statistically indistinguishable from untreated oocysts. This provides direct evidence that the outer layer is expendable, contributing little to overall structural integrity and mechanics of the oocyst wall, whereas the inner layer is effectively self-sufficient as a mechanical barrier. This interpretation is reinforced by concordant AFM data from T. gondii, demonstrating that the inner layer alone exhibits mechanical robustness equivalent to that of the complete bilayered oocyst wall ^17^. At the molecular level, this resilience is likely supported by the β-1,3-glucan scaffold described in the inner layer of *T. gondii* and *E. tenella* oocyst walls ^18^. This scaffold forms a load-bearing fibrillar network that, together with dityrosine cross-links localised within the inner wall layer, is proposed to provide structural rigidity ^14,16^.

The lectin permeability data extend this hierarchy from mechanics to molecular transport, while also providing molecular markers that distinguish the different wall layers. Bleach- and ozone-treated oocysts remain impermeable to the PNA, a ∼110-kDa macromolecule, supporting the view that the inner wall constitutes a highly selective permeability barrier inaccessible to macromolecules from the exterior even after outer-layer loss. Conversely, the marked increase in WGA and MPL binding after bleach treatment indicates that the intact outer layer physically occludes glycan motifs on the inner wall surface, which become accessible only upon its removal. Ozone produced a qualitatively distinct wall state: although it similarly disrupts the outer layer, WGA and MPL signals remained low, comparable to those observed in untreated oocysts, suggesting that ozone simultaneously removes the outer layer and oxidises the exposed glycans in the inner layer. This effect is more pronounced for WGA-reactive motifs, which appear more susceptible to oxidative degradation than MPL-reactive Gal/GalNAc residues. Despite both eroding the outer layer, bleach and ozone produce functionally distinct wall states; under our experimental conditions, ozone was more aggressive than bleach and extended its activity to glycan components of the inner wall.

The impermeability of this inner layer also has a mechanical corollary: a wall that excludes macromolecules and limits exchange can sustain an internal pressure that does not equilibrate with the environment, defining the oocyst’s behaviour as a sealed, pressurised shell. To test whether the external environment can modulate this pressure, we challenged oocysts across osmotic conditions spanning hypo-osmotic freshwater, near-isosmotic physiological buffer, and the strongly hyperosmotic regimes (10× PBS) that exceed the osmolarity of seawater, probing the upper bound of osmotic stress that oocysts could encounter in saline or evaporating environments ^8^.

Across this range, osmotic perturbations produced only small changes in estimated internal pressure and in rupture force, far smaller than the collapse expected if internal pressure equilibrated passively with the environment. This indicates that the wall behaves as a largely sealed, pressure-bearing shell rather than a passively equilibrating membrane. Our data further separate two mechanical regimes: in the pre-rupture elastic regime, indentation stiffness is well described by a pressure-dominated shell model, so internal pressure contributes substantially to resistance to deformation; yet rupture force was neither reduced by osmotic challenge nor by outer-layer removal, both of which would be expected if pressure governed failure. Ultimate wall failure is therefore set by the intrinsic material properties of the inner layer, independently of internal pressure.

Such functional partitioning between wall layers is a recurring principle in walled biological systems. In fungal and plant cells, distinct polysaccharide networks fulfil distinct mechanical and chemical roles, with load-bearing scaffolds, including β-glucans and chitin in fungi, cellulose in plants, spatially separated from surface-modulating components such as mannans and pectins ^19^. A key difference, however, is that plant and fungal cells maintain turgor through regulated water and solute exchange across their wall to drive cell expansion and growth. The coccidian oocyst wall, by contrast, appears to behave as a constitutively sealed protective barrier, limiting passive solute exchange and functioning mechanically as a closed pressure-bearing shell. Within this context, the β-1,3-glucan and dityrosine-crosslinked inner layer fulfils an analogous structural role to load-bearing polysaccharide networks in fungi and plants, while the outer layer primarily modulates surface chemistry and environmental interactions ^15^.

The second determinant is that this globally resistant wall contains a spatially defined mechanical weak point. The anterior pole ruptured at lower forces than lateral or posterior regions, and sporocysts were frequently ejected through this site even when force was applied elsewhere. These observations indicate that the anterior pole functions as a mechanically privileged exit gate, a spatially fixed weak point that directs sporocyst release under force, providing a mechanical basis for a spatially regulated excystation architecture in which polar-region weakness is intrinsic to the wall and sufficient to direct release of the sporocysts. This mechanical anisotropy helps explain the sequential logic of excystation in avian *Eimeria in vivo*. Excystation proceeds through two successive steps: mechanical rupture of the oocyst wall in the gizzard releases sporocysts, after which trypsin and bile salts induce sporozoite release from the sporocysts ^30^. *In vitro* protocols recapitulate this sequence by first mechanically disrupting oocysts, typically with glass beads ^31^, before exposure to digestive factors. Consistent with this model, trypsin and/or bile salts alone were insufficient to open intact oocysts, trigger sporozoite release, or measurably weaken wall mechanics, whereas mechanically opened oocysts underwent efficient excystation after exposure to digestive factors, as previously described ^30^. These data support a model in which the oocyst wall must therefore be mechanically breached *in vivo* by relatively high forces, before digestive factors can act on the sporocyst wall, a requirement physiologically met by the grinding action of the gizzard, which oocysts traverse before encountering bile and pancreatic enzymes in the intestine.

How the coccidian oocyst wall is breached in non-avian hosts in the absence of a dedicated mechanical grinding organ remains an open question. In mammalian *Eimeria* species, specialized wall-associated structures, including caps or regions of reduced thickness such as the micropyle, are thought to represent preferential sites for a simultaneous chemical attack by reducing agents, e.g., cysteine-HCl, CO₂, and/or proteolytic enzymes such as pepsin ^2,12,32^. For coccidian parasites lacking such structures, including *C. cayetanensis* and *T. gondii*, the mechanisms governing sporocyst release during infection remain poorly understood and may involve biochemical weakening of the wall within the gastrointestinal tract and/or mechanical stresses generated by intestinal peristalsis, which together may contribute to wall rupture and subsequent parasite release.

The structural basis of anterior-pole weakness remains to be determined. One contributor is the *Eimeria* oocyst’s own geometry: the anterior and posterior poles differ in shape, with the anterior pole being more sharply curved, and such local curvature differences could influence how mechanical stress is distributed under load. Additional factors may also contribute, including local differences in wall thickness, molecular composition, cross-link density, glucan scaffold continuity, or the presence of a micropyle. The rupture forces recorded here are orders of magnitude higher than those required to puncture a cellular membrane by AFM (∼10–100 nN) ^33,34^, but broadly comparable in magnitude to the bursting forces of other walled microorganisms measured by micromanipulation, such as baker’s yeast ^35^, placing the coccidian oocyst wall within a broad class of biological barriers that combine high global resistance with local mechanical specialisation. The coccidian oocyst appears to use an analogous spatial biomechanical logic, tuned for transmission rather than growth: a globally resistant wall protects the infectious sporozoites in the environment, while a mechanically privileged anterior pole facilitates their release when the appropriate physical and biochemical cues are encountered in the host.

The third determinant is that structural or chemical alterations of the wall are decoupled from the infectivity, as demonstrated here in *E. tenella* and previously in *E. acervulina* and *T. gondii* ^24^. In our experimental conditions, bleach eroded the outer layer entirely and strongly altered wall surface accessibility yet left the parasite fully infective, whereas heat, which spared both the bilayer and its surface glycans, abolished infectivity outright. Thus, visible wall damage does not necessarily predict loss of parasite viability. Conversely, preservation of wall ultrastructure does not guarantee infectivity. Autofluorescence, used to report structural and molecular changes in the oocyst wall, reflects neither mechanical properties nor infectivity. The dissociation between mechanics and ultrastructure suggests that thermal damage acts at the molecular level, likely through protein denaturation and aggregation, and reorganisation or disruption of dityrosine cross-links within wall protein networks, leading to increased material heterogeneity and reduced resistance to rupture, as reported in other complex polymer networks ^36,37^. The drop in estimated internal pressure in ozone- and heat-treated oocysts may reflect denaturation or aggregation of internal contents.

The two species also registered treatment-induced damage differently. In *E. acervulina*, heat and ozone reduced median rupture force, indicating direct weakening of the load-bearing inner layer. In *E. tenella*, the same treatments left median rupture force unchanged and instead reduced internal pressure while increasing the proportion of mechanically weakened oocysts. This suggests that *E. tenella* has a more mechanically resilient inner wall under the conditions tested, possibly reflecting differences in molecular cross-linking or scaffold organisation. Notably, our findings align with evidence obtained in other coccidian models, supporting the view that environmental resistance differs among chicken *Eimeria* species and more broadly among coccidian parasites, although the specific pattern of resistance depends on the external stressor considered^11^.

Functional differences between oocyst wall layers have important implications for controlling coccidian oocyst transmission using existing disinfection strategies. Ozone is a powerful disinfectant, widely used in drinking water treatment and proposed as a food-compatible alternative to chlorination for decontaminating ready-to-eat fresh produce ^38^. Our data indicate that treatment with 20 mg/L ozone, ten times the dose typically used in industrial settings, induces wall alterations comparable to bleach, with both producing the largest decrease in autofluorescence. Ozone also weakens the wall more than bleach, but less than heat. However, it does not achieve complete sporozoite inactivation at this dose and exposure time. Because the inner wall governs both the mechanical resistance and the impermeability of the oocysts to chemical agents, reliable inactivation requires breaching the inner wall and likely the sporocyst wall, or direct access to the sporozoites. In practice, few of the treatments currently used in the food and water industries meet these requirements. They include high-pressure processing and high doses of gamma or ultraviolet irradiation, though their efficacy against coccidian oocysts may further vary depending on the composition of the matrix being decontaminated ^38^.

In conclusion, this work establishes that the coccidian oocyst wall is not a homogeneous barrier but an architecturally specialised structure in which two functionally distinct layers underpin environmental resistance, such as permeability, mechanically gated excystation and spatial regulation of sporocyst release. These determinants are likely to extend beyond *Eimeria* to other coccidians, including *T. gondii* and potentially *C. cayetanensis*, which are parasites of major public health significance due to their widespread distribution and their impact on human health, given the conserved bilayered organisation and shared capacity to withstand various environmental conditions including chemical disinfectants ^3,14,17^. Future work should determine whether anterior-pole weakness is attributable to specific protein distributions, local wall thickness or discontinuities in the β-glucan scaffold, and whether targeted disruption of inner wall components, particularly at these mechanically weaker zones, can serve as the basis for new decontamination or anti-infective strategies.

## Methods

### Ethics statements

All animal procedures, including *in vivo* experiments and *Eimeria* oocyst production, were conducted in accordance with EU Directive 2010/63/EU and French regulations on animal experimentation (Rural Code 2018; Decree No. 2013-118). Experiments at the PFIE facility, INRAE Tours-Nouzilly were approved by the CEEAVdL ethics committee (Committee No. 19; APAFIS No. 25884). Oocyst production at ANSES was approved by Ethics Committee No. 16 (APAFIS No. 34204).

### Oocyst production

*Eimeria acervulina* oocysts were produced at two independent sources, INRAE Nouzilly and ANSES Ploufragan, following the regulatory-mandated closure of animal facilities at INRAE Nouzilly. *Eimeria tenella* oocysts were produced exclusively at INRAE Nouzilly. Irrespective of the *Eimeria* species or production site, all oocyst batches were received after sporulation, shipped at 2–8 °C, and stored at 4 °C in 2% (w/v) aqueous potassium dichromate, K2Cr2O7. *Eimeria* oocyst batches were considered heterogeneous biological populations, as individual oocysts may differ in maturity, sporulation status, physiological state and infectivity. Oocysts were used within six months of production, as infectivity declines progressively thereafter ^39^.

### *E. acervulina* INRAE production

White Leghorn chickens (4-6 weeks old) were orally inoculated with 10⁵ sporulated oocysts. Faecal samples were collected at 5–6 days post-inoculation (dpi), with six birds per cage. Faeces were filtered through a series of sieves with decreasing mesh sizes (1000–100 µm), and oocysts were concentrated by centrifugation at 1050 × *g* for 5 min at room temperature (RT). Oocysts were purified by magnesium sulphate (MgSO4) flotation (specific gravity 1.20) at 1050 × *g* for 6 min at 18 °C. To limit bacterial contamination, oocysts were incubated for 20 min at RT in an aqueous solution containing 0.35% active chlorine, obtained by a 1:10 dilution of 3.5% NaOCl solution (VWR Chemicals). Oocysts were subsequently washed three times in distilled water by centrifugation at 1050 × *g* for 6 min. Sporulation was performed at 26 °C under agitation in 2% K2Cr2O7 until at least 95% of oocysts were sporulated. Sporulated oocysts were enumerated using a Thoma chamber and stored at 4 °C in 2% K2Cr2O7.

### *E. acervulina* ANSES production

ISA Brown chickens (21–35 days old) were orally inoculated with 5 × 10⁴ sporulated oocysts in 1 mL sterile saline. Faecal droppings were collected at 4–6 dpi, and oocysts were purified by sodium chloride flotation (specific gravity 1.18) ^40^. Sporulation was performed in 2% K2Cr2O7 on agar plates at 28 °C and 65% relative humidity for 24 h. Sporulated oocysts were washed, enumerated by Neubauer haemocytometer, and stored in 2% K2Cr2O7 at 4 °C ^24^.

### *E. tenella* INRAE production

White Leghorn chickens (4–6 weeks) were orally inoculated with 10⁴ sporulated *E. tenella* oocysts and maintained under coccidian-free conditions. At 7 dpi, caeca were collected and scraped. Scrapings and caecal content were homogenised for 1 min using an autoclavable steel blender. Oocysts were recovered by centrifugation at 750 × *g* for 5 min at RT. To avoid aggregation, the oocyst suspension was dissociated by incubation in PBS containing 1.5% (w/v) trypsin (Sigma-Aldrich, T4799) for 45–60 min at 41 °C. Oocysts were then purified by MgSO4 flotation (specific gravity 1.20) at 750 × *g* for 5 min. To limit bacterial proliferation, oocysts were incubated with 0.35% active chlorine for 20 min at RT prior to being washed three times in distilled water. Sporulation was performed at 27 °C in 2% K2Cr2O7 for 48–72 h under continuous agitation. Sporulated oocysts were enumerated by Thoma haemocytometer and stored at 4 °C in 2% K2Cr2O7.

### Oocyst treatments

Prior to all treatments, oocysts were washed three times in PBS (pH 7.4; Gibco, cat. no. 14190-094) by centrifugation at 900 × *g* for 5 min at RT to remove residual K₂Cr₂O₇. A suspension of 10⁶ oocysts in 1 mL PBS was used per condition unless stated otherwise.

### Digestive factors

An *in vitro* excystation medium was used to mimic digestive conditions associated with *Eimeria* oocyst excystation *in vivo* ^41^. Oocysts were incubated in PBS containing 0.4% (w/v) trypsin (Sigma-Aldrich, T4799), and 1.6% (w/v) bile salts (Sigma-Aldrich, B8631). The pH was adjusted to 7.4–7.6 with 1.0 N NaOH (Alfa Aesar, cat. no. 35629). Incubation was performed for 10 min at 41 °C under static conditions. In parallel, oocysts were treated with trypsin alone, bile salts alone, or PBS alone as a control. After incubation, oocysts were pelleted by centrifugation at 900 × *g* for 5 min and washed once in PBS.

### Disinfection treatments

For bleach treatment, oocysts were incubated in 1 mL PBS containing 3% active chlorine, prepared from a commercial bleach solution (Lacroix Javel, Colgate-Palmolive) for 30 min at 4 °C. This treatment has been reported to selectively remove the outer oocyst wall of *T. gondii* without abolishing infectivity ^17^. Oocysts were then washed three times in PBS by centrifugation at 900 × *g* for 5 min and resuspended in 500 µL PBS.

Heating oocysts consisted in incubating the parasites in 1 mL PBS at 80 °C for 10 min in a calibrated dry block heater, rendering them non-infective ^17^. Suspension osmolarity was verified before and after heating by Mikro-Osmometer (Löser Messtechnik); no change was observed. Samples were cooled passively to RT before analysis.

Ozone experiments were performed using a pilot-scale ozone system. Ozone was dissolved in 1 L autoclaved demineralised water at 15 °C in a 3 L stainless-steel tank, with continuous amperometric monitoring (Q46H64 probe, Équipements Scientifiques). Oocysts were exposed to a target ozone concentration of 20 mg/L for 60 min at RT. Two independent *E. tenella* assays were performed using 1.8 × 10⁷ and 1.9 × 10⁷ oocysts, respectively, and one *E. acervulina* assay was performed using 5.8 × 10⁷ oocysts. Mean measured ozone concentrations were 19.7 ± 0.58 and 24.1 ± 3.90 mg/L for the two *E. tenella* assays, and 21.4 ± 2.50 mg/L for the *E. acervulina* assay, expressed as mean ± s.d. After exposure, residual ozone was eliminated by air bubbling for 12, 40, and 20 min, respectively. Oocysts were then recovered by filtration through 0.45 µm cellulose ester membranes (Millipore), eluted with PBS, and concentrated by centrifugation at 3500 × *g* for 15 min at 10 °C.

### Osmotic shocks

Oocysts were pelleted by centrifugation at 900 × *g* for 5 min from PBS (286 mOsm/L). Two parallel conditions were established simultaneously. For hyperosmotic exposure, oocysts were incubated in 1 mL 10× PBS (2817 mOsm/L; Gibco, cat. no. 14200-067) for 24 h at 4 °C. Isotonic conditions were then restored by diluting 100 µL of the hyperosmotic suspension with 900 µL ultrapure water, yielding a final osmolarity of 282 mOsm/L, confirmed using a Mikro-Osmometer (Löser Messtechnik), and incubated for a further 24 h at 4 °C. For hypoosmotic exposure, a separate oocyst aliquot was incubated in 1 mL ultrapure water (∼0 mOsm/L) for 24 h at 4 °C. After incubation, all samples were centrifuged, resuspended in PBS, and processed immediately for morphological and mechanical analyses.

### Fluorescence labelling, flow cytometry, and microscopy

For all fluorescence assays, oocysts were washed three times in PBS by centrifugation at 900 × *g* for 5 min at RT and adjusted to 2 × 10⁶ oocysts/mL.

### Lectin-binding assay

TRITC-conjugated lectins, wheat germ agglutinin WGA, *Maclura pomifera* lectin MPL and peanut agglutinin PNA were selected based on their reported binding to *Eimeria* and *Toxoplasma* oocyst walls ^24–26^. These lectins were used as wall-binding probes rather than as definitive markers of specific oocyst wall glycans, targeting N-acetylglucosamine/sialic acid, galactose-containing structures, and β-galactose, respectively. Fluorescein (FITC)-conjugated lectins were used but exhibited a large spectral overlap with oocyst wall autofluorescence, impeding their use for reliable quantification.

Oocysts were incubated with TRITC-WGA (Vector Laboratories, RL-1022), TRITC-MPL (bioWORLD, 21761039-1), TRITC-PNA (bioWORLD, 21761000-1) at 50 µg/mL in 100 µL PBS for 2 h at RT in the dark, washed once in PBS by centrifugation at 900 × *g* for 5 min and resuspended in 400 µL PBS. Unlabelled controls, consisting of oocysts incubated without lectin, were included as negative controls for each condition. For positive controls, oocysts were pre-incubated with 0.1% (w/v) poly-L-lysine (PLL; Sigma-Aldrich, P8920) for 60 min prior to lectin labelling for 2 h under the same conditions. PLL, a polycationic polymer, was used empirically as a positive control to enhance surface-associated lectin labelling.

### Flow cytometry

Measurements were performed on a BD LSRFortessa X-20 (BD Biosciences) equipped with 561, 488, 640, and 405 nm lasers. TRITC fluorescence was excited at 561 nm and detected using a 586/15 nm bandpass filter. Detector voltages were optimised at the beginning of the experiment and kept constant for all acquisitions. Instrument performance was verified using CS&T beads (BD Biosciences).

Oocysts were identified using FSC-A and SSC-A parameters, and doublets were excluded using FSC-A versus FSC-H gating. At least 10⁴ events were acquired per sample. Each condition was analysed in five independent experiments. Data were analysed using FlowJo v10.10.0 (BD Biosciences). Lectin binding was expressed as the fold change in geometric mean fluorescence intensity (gMFI), calculated as:

Fold change = gMFI-labelled / gMFI-unlabelled

where the unlabelled control corresponded to the same experimental condition.

### Fluorescence microscopy

Lectin labelling was visualised on an Axio Observer.Z1 inverted microscope (Zeiss) with a Plan-Apochromat 40×/1.3 NA oil objective, CoolLED pE-400max at 100% diode intensity, Zeiss filter set 20 HE (excitation 546/12 nm, emission 575–640 nm), ORCA-Flash4.0 LT Plus camera (Hamamatsu) and motorized stage (Märzhäuser Wetzlar). Images were acquired using Micro-Manager v1.4.22 ^42^ with an exposure time of 300 ms. Microscopy images were used for qualitative visualization of lectin labelling; quantitative analysis was performed by flow cytometry.

### Autofluorescence imaging

Images for oocyst segmentation and wall autofluorescence quantification were acquired on an Axiovert 200M inverted microscope (Zeiss) with 40×/0.9 NA air objective, CoolLED pE-300max, quadriband filter (F66-888, AHF Analysentechnik) and CoolSnap HQ2 camera (Photometrics). Images were acquired using Micro-Manager v1.4.22 ^42^ with excitation at 365 nm. A uniform display threshold was applied for figure presentation; quantification used unthresholded raw images.

### Image analysis and oocyst segmentation

Oocysts were segmented using a StarDist2D deep learning pipeline ^43^ implemented in Python with the Stardist/CSBDeep framework ^44^. Training annotations were generated in Labkit within Fiji/ImageJ ^45^, with labels corresponding to sporulated oocysts. Images were percentile-normalised and split into 80:20 training and validation datasets with a fixed random seed. The StarDist2D model used a U-Net backbone, 32 radial rays and an 8 × 8 grid. Probability and non-maximum-suppression thresholds were optimised on the validation dataset. The trained model is available as described in Code Availability.

Segmented objects were quantified in Celldetective v 1.4.1 ^46^. Oocyst area was calculated from calibrated images, using a pixel size of 0.16 µm per pixel for Nikon TiE inverted microscope. For each frame, the mean background intensity was measured and subtracted from all pixel values prior to quantification. Wall autofluorescence was quantified as the mean intensity of pixels located within 6 pixels of the segmented contour, determined by a Euclidean distance transform. Measurements were computed with scikit-image within Celldetective.

### Transmission electron microscopy

Oocysts suspended in 100 µL PBS were partially disrupted by pulsed sonication to facilitate resin infiltration during embedding. Sonication was performed using a Vibra-Cell sonicator (20 kHz, 60% amplitude) with an Arduino-controlled triggering device, delivering reproducible 1 s pulses (15–45 min on ice).

Samples were fixed in 2.5% v/v glutaraldehyde (Sigma-Aldrich, G5882) in 200 mM HEPES buffer (pH 7.2; Thermo Fisher, 15630056), washed in HEPES buffer, concentrated in 2% low-melting-point agarose (Sigma-Aldrich, A9414), and post-fixed in 1% osmium tetroxide (EMS, 19150) for 1 h at 4 °C. Samples were then dehydrated through a graded ethanol series followed by propylene oxide and embedded in Spurr’s resin. Ultrathin sections of 60–90 nm were stained with uranyl acetate and lead citrate and examined using a Tecnai 200 kV TEM (Thermo Fisher Scientific) equipped with a 4K × 4K OneView camera (AMETEK). Image exposure was set automatically; therefore, grey-level intensities were not compared quantitatively between conditions.

### Mechanical characterisation

#### Microindentation

Single oocyst mechanics were measured by micromanipulation-based microindentation as described ^28^. Oocysts were held by hydrostatic aspiration (∼50 Pa, equivalent to ∼5 mm H₂O) using a holding micropipette with a tip diameter of 7–8 µm for *E. acervulina* and 8–9 µm for *E. tenella*.

Micropipettes were pulled from borosilicate capillaries (Harvard Apparatus) using a P-97 puller (Sutter Instruments) and calibrated against reference probes (Aurora Scientific, model 406A). The holding pipette was mounted on a piezoelectric stage (Physik Instrumente) on a motorised micromanipulator (MP-285, Sutter Instruments).

Oocysts were indented laterally using externally calibrated flexible micropipettes ^47^. Needle-shaped probes with stiffnesses of 7200 ± 250 or 9730 ± 1230 nN/µm were used.

Probe deflection was tracked optically at ∼400 Hz with a sub-pixel accuracy of 20–30 nm on a Nikon TiE inverted microscope mounted on an air-suspension table. Images were acquired simultaneously at 1 frame/s during indentation assays using Micro-Manager v1.4.22 and a custom MATLAB R2021b routine. Force was calculated as *F* = *k_indent_δ*, where *k_indent_* is the calibrated probe stiffness (nN/µm) and *δ* is the measured probe deflection (µm). Rupture force was defined as the maximum force reached before abrupt wall failure.

Thirty oocysts were analysed per condition, with a matched untreated control included in each experimental session; exclusion criteria included focus loss, detachment, or ambiguous contact. Data were processed in Python 3.10.18 using the analysis pipeline described in Code Availability. A representative indentation is shown in Supplementary Video 1.

#### Internal pressure estimation

Internal pressure was estimated from microindentation stiffness using the large-indentation framework of Vella et al. ^29^, in which:

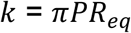

where *k* is the contact stiffness, i.e. the slope of the force versus indentation, *P* is the internal pressure and *R_eq_* is the equivalent local radius of curvature at the indentation site.

For non-spherical oocysts, *R_eq_* is defined by 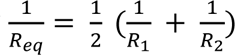, i.e. its inverse equals the mean of the two principal curvatures.

Contact stiffness was extracted as the slope of force–indentation curves over the linear 1–5 µm indentation range using `scipy.stats.linregress`. Local radii of curvature were measured from brightfield images by fitting circles to oocyst contours at each indentation site using custom Python code. Top and bottom regions were approximated as spherical caps (*R*_1_ = *R*_2_), whereas side regions were treated as ovoidal surfaces with two principal radii. Radius measurements were matched to stiffness measurements using oocyst identifiers.

For each condition, pressure was estimated by fitting a linear model through the origin to paired *R_eq_* and *k* values:

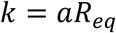

where the fitted slope corresponds to *πP*. Pressure was therefore calculated as:

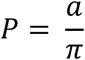

Pressure values derived from this fit are expressed in kPa (1 nN/µm² = 1 kPa). The uncertainty of *P* was derived from the standard error of the fitted slope. The coefficient of determination, *R*^2^, was calculated to assess goodness of fit. To assess whether the pressure-dominated model was consistent across oocyst geometry, *k* was plotted as a function of *R_eq_* for anterior, lateral and posterior measurements. Under the pressure-dominated model, measurements from all orientations are expected to collapse onto a common line through the origin.

#### Atomic force microscopy

For AFM measurements, oocysts were immobilised either on plasma-cleaned glass-bottom dishes coated with 0.01% (w/v) poly-L-lysine or by confinement within custom PDMS microstructured wells when adhesive immobilisation was insufficient. Measurements were performed on a NanoWizard 4XP AFM (JPK/Bruker) mounted on a Zeiss Axiovert 200 inverted microscope and operated in closed-loop mode at RT.

Silicon nitride cantilevers (MLCT-BIO-DC, Bruker; nominal spring constant 10–30 pN/nm) were calibrated before each experiment by determining the deflection sensitivity on glass and the spring constant using the thermal noise method. Force–distance curves were acquired in PBS at RT with a ramp speed of 2 µm/s, a 7 µm z-ramp and a 1 nN force setpoint. Up to five curves were acquired per oocyst.

Curves were processed using JPK Data Processing software. After baseline correction and contact point determination, the approach branch was fitted with the Hertz–Sneddon model for a quadratic pyramidal indenter, assuming an incompressible material with *ν* = 0.5. Fits were restricted to the initial indentation region to minimise contributions from deeper structures. For each oocyst, a mean apparent Young’s modulus was calculated from force curves. When relevant, adhesion forces were extracted from the retraction branch.

#### Infectivity assays

To assess the effect of disinfectant treatments on parasite infectivity, *E. tenella* oocysts were subjected to the indicated treatments before oral inoculation of White Leghorn chickens. In total, 11 independent groups of 20 chickens were infected with 10⁴ sporulated oocysts per chicken, corresponding to three independent experimental replicates for the untreated control, bleach, and ozone conditions, and two independent replicates for the heat condition. Two replicates were performed for the heat condition as complete abolition of infectivity, evidenced by the total absence of caecal lesions and undetectable oocyst output in both replicates, was determined to preclude the need for a third independent experiment. At 7 days post-infection, chickens were euthanised, oocysts recovered from the caeca were counted, and caecal lesions were scored on a scale from 0 to 4 according to Johnson and Reid scoring system ^48^, as a semi-quantitative readout of *E. tenella* pathogenicity, where 0 indicates no lesions and 1–4 reflect increasing severity of caecal pathology from mild petechial haemorrhage to extensive tissue damage and mortality.

#### Statistics

Statistical analyses were performed in Python using NumPy, pandas and SciPy, and figures were generated using matplotlib and seaborn. Given the broad distributions of the data, groups were compared using two-tailed Mann–Whitney U-tests throughout. Data are presented as median with interquartile range unless stated otherwise. Statistical significance was defined as *P* < 0.05. Effect sizes were quantified using Cliff’s delta (δ) ^49^, a non-parametric measure, computed for each treatment relative to the corresponding untreated control. δ was classified as negligible (|δ| < 0.147), small (0.147 ≤ |δ| < 0.33), medium (0.33 ≤ |δ| < 0.474) or large (|δ| ≥ 0.474), following Romano et al ^50^. Effect sizes for all comparisons are reported in Supplementary Table 1. Replicate numbers are indicated in figure legends.

## Acknowledgements

This work was supported by the ANR project BreakingTheWall (ANR-22-CE35-0008), which also funded J.E.H.’s PhD. The IMM microscopy core facility is member of the national infrastructure France-BioImaging supported by the French National Research Agency (ANR-24-INBS-0005 FBI BIOGEN). We thank Martine Pelicot and Evelyne Thi Tien Nguyen (LAI, Aix Marseille Université) for assistance with flow cytometry, and Philippe Robert (LAI, Aix Marseille Université) for designing and building the Arduino-controlled triggering device used for sonication experiments. The authors are grateful to the members of the scientific and animal staff of the Plateforme d’Infectiologie Expérimentale (PFIE, INRAE, 2026. Infectiology of farm, model and wild animal’s facility, https://doi.org/10.15454/1.5572352821559333e12), UE-1277 PFIE, INRAE Centre Val de Loire, Nouzilly, France, especially to Experimental Zone team, in particular Thierry Chaumeil, Olivier Dubès and Lorine Branger. PFIE is part of EMERG’IN, the national infrastructure for the control of animal and zoonotic emerging infectious diseases through *in vivo* investigation.

## Author contributions

J.E.H. performed microindentation, AFM, autofluorescence imaging, lectin labelling, flow cytometry experiments, deep-learning segmentation and data analysis, and wrote the original draft.

R.T. developed the Celldetective analysis framework used for image segmentation and measurement extraction.

L.S. performed the *in vivo* infectivity assays and contributed to *E. tenella* oocyst production.

C.C. performed the ozone treatment experiments.

H.L.G. and A.K. performed transmission electron microscopy sample preparation and imaging.

J.-M.R. produced *E. acervulina* oocysts at ANSES.

M.R. supervised the chicken breeding, ethical process (*in vivo* part) and infection facilities in confined medium.

S.L.C. supervised the ozone treatment experiments and provided infrastructure.

A.S. supervised the *in vivo* infectivity assays, *E. tenella* oocyst production, and analysis of *in vivo* infectivity data.

J.H. conceived and designed the study, supervised experimental work, developed the microindentation framework and internal pressure estimation model, contributed to data interpretation, acquired funding and wrote the manuscript with input from all authors.

P.-H.P. conceived and designed the study, supervised experimental work, contributed to the design of the microindentation and AFM experiments, supervised the mechanical characterisation, acquired funding and wrote the manuscript with input from all authors.

A.D. conceived and designed the study, supervised all experimental work, acquired funding, and wrote the manuscript with input from all authors.

All authors reviewed and approved the final manuscript. J.H., P.-H.P. and A.D. jointly supervised this work.

## Data availability

The data supporting the findings of this study are available within the article, its Supplementary Information and Source Data files. Additional data are available from the corresponding authors upon request.

## Code availability

The microindentation analysis pipeline used in this study was described previously ^28^ and is available on Zenodo (https://doi.org/10.5281/zenodo.17467539). The oocyst segmentation pipeline, including the StarDist model trained in this study, is available on Zenodo (https://doi.org/10.5281/zenodo.20732270). Single-cell measurements were extracted using Celldetective ^46^, an open-source package available at https://github.com/celldetective/celldetective.

## Supplementary Information

**Supplementary Fig. 1.**
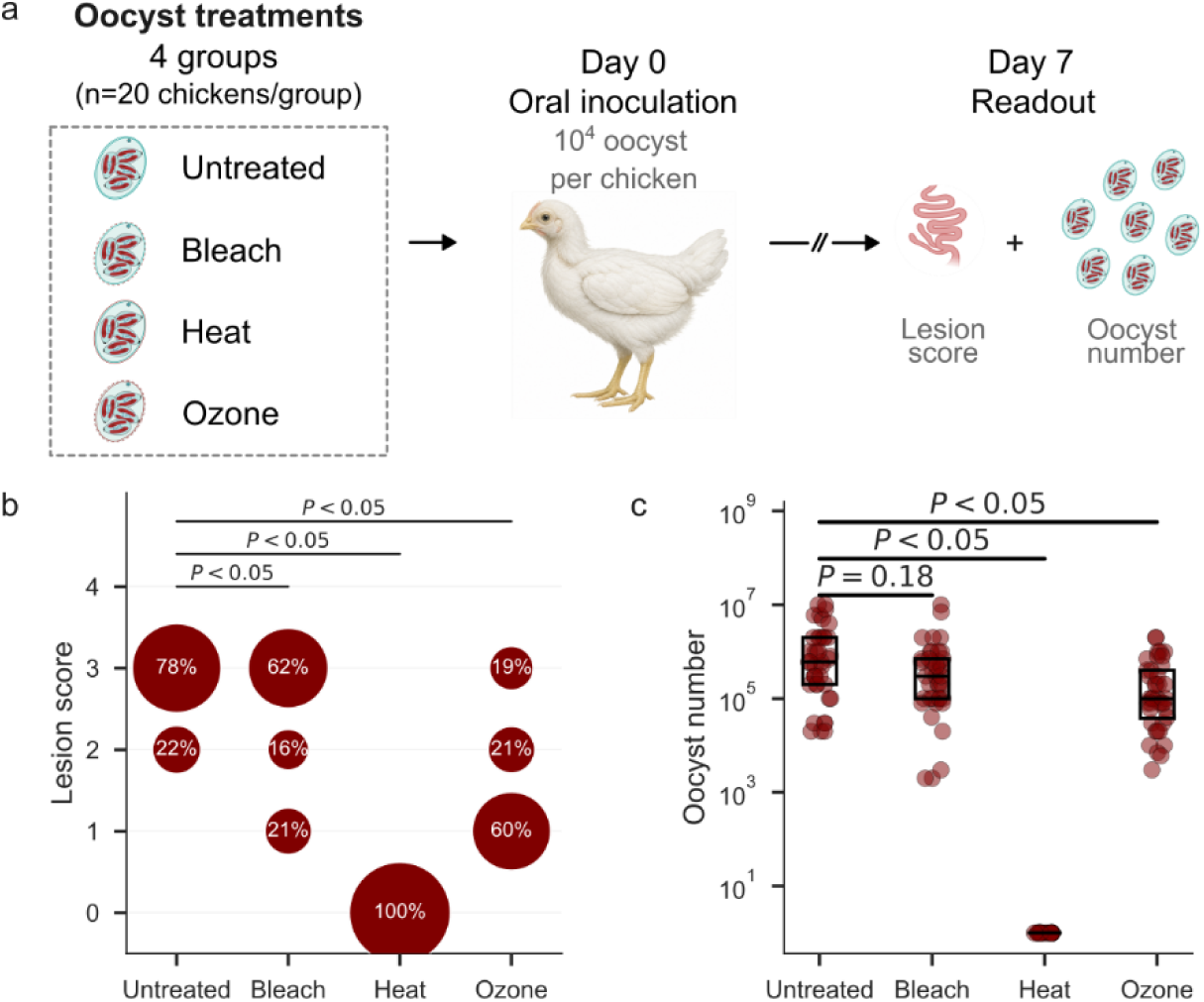
| Heat treatment reduces the infectivity of *Eimeria* oocysts. **a**, Experimental design. Oocysts were left untreated or exposed to bleach, heat, or ozone before oral inoculation of chickens on day 0. Each chicken received 10⁴ oocysts, with four groups of 20 chickens per group (three replicates for untreated, bleach, and ozone; two replicates for heat, represented here as pooled data per treatment condition). Lesion scores and faecal oocyst counts were assessed at day 7 post-inoculation. **b**, Caecal lesion scores at day 7 post-inoculation with untreated or treated oocysts, assessed on a scale of 0–4, where higher scores reflect increasing severity of caecal pathology. **c**, Faecal oocyst numbers measured at day 7 after inoculation with untreated or treated oocysts. In **b,c**, each dot represents an individual chicken; boxes indicate the interquartile range and horizontal lines indicate the median. *P* values were calculated using two-sided Mann–Whitney *U* tests for the indicated pairwise comparisons.

**Supplementary Fig. 2.**
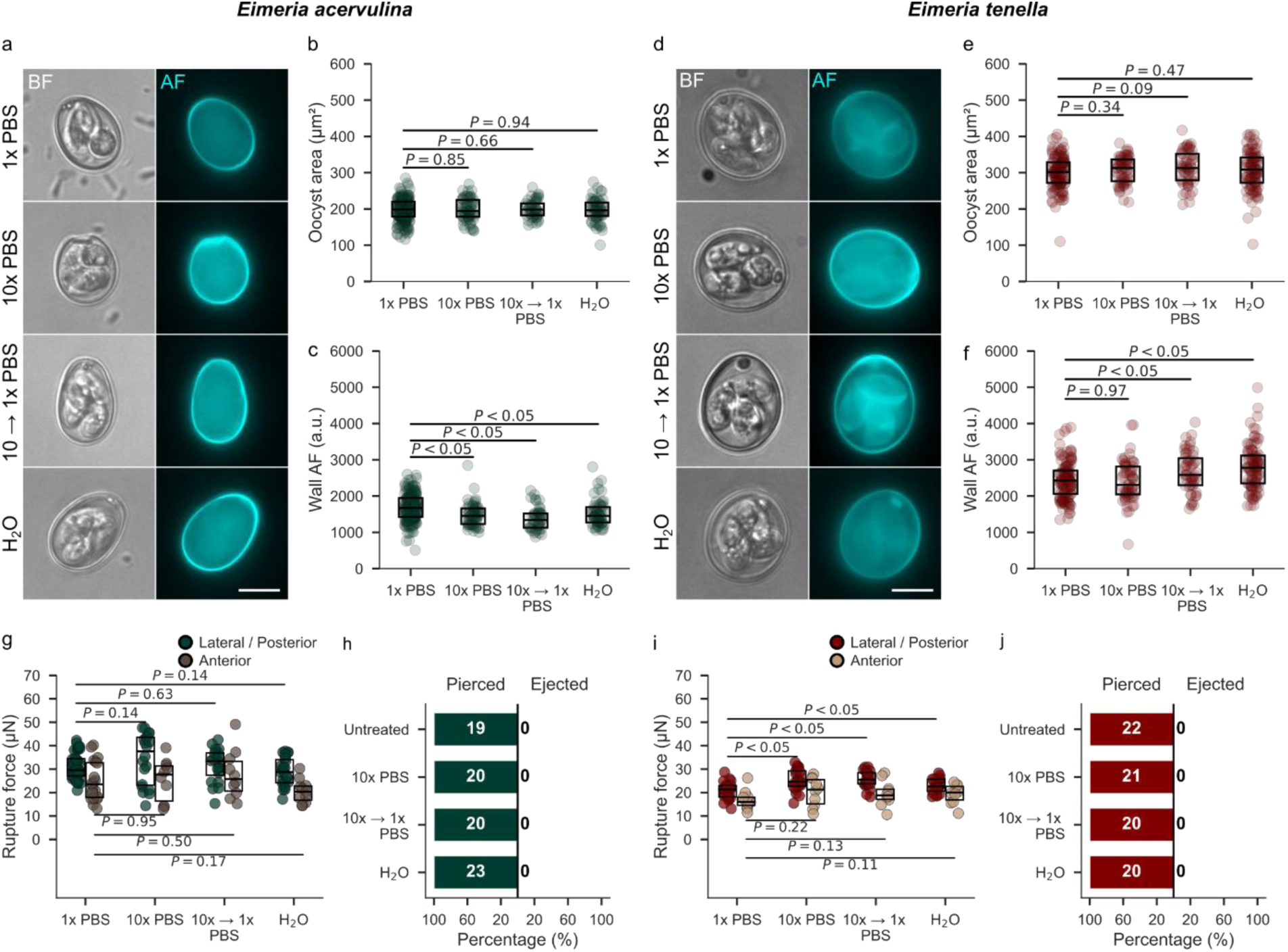
| Osmotic perturbations modestly affect oocyst wall autofluorescence and rupture force in *Eimeria acervulina* and *E. tenella*. **a**, Representative bright-field (BF) and autofluorescence (AF) images of *E. acervulina* oocysts under 1× PBS, 10× PBS, 10× PBS → 1× PBS, and H₂O conditions for 24 h. Scale bars, 10 µm. **b**, Quantification of oocyst projected area in *E. acervulina* across osmotic conditions. **c**, Quantification of oocyst wall autofluorescence intensity in *E. acervulina*. a.u., arbitrary units. **d–f**, Same as **a–c** for *E. tenella*. **g**, Rupture force of *E. acervulina* oocysts under osmotic conditions. Oocysts were indented at the lateral/posterior position or at the anterior pole, as indicated by the colour code. **h**, Percentage of *E. acervulina* oocysts that were pierced or showed content ejection during microindentation under osmotic conditions. Numbers inside bars indicate the number of oocysts in each category. **i,j**, Same as **g,h** for *E. tenella*. Each dot represents an individual oocyst; boxes indicate the interquartile range and horizontal lines indicate the median. *P* values were calculated using two-sided Mann–Whitney *U* tests for the indicated pairwise comparisons.

**Supplementary Fig. 3.**
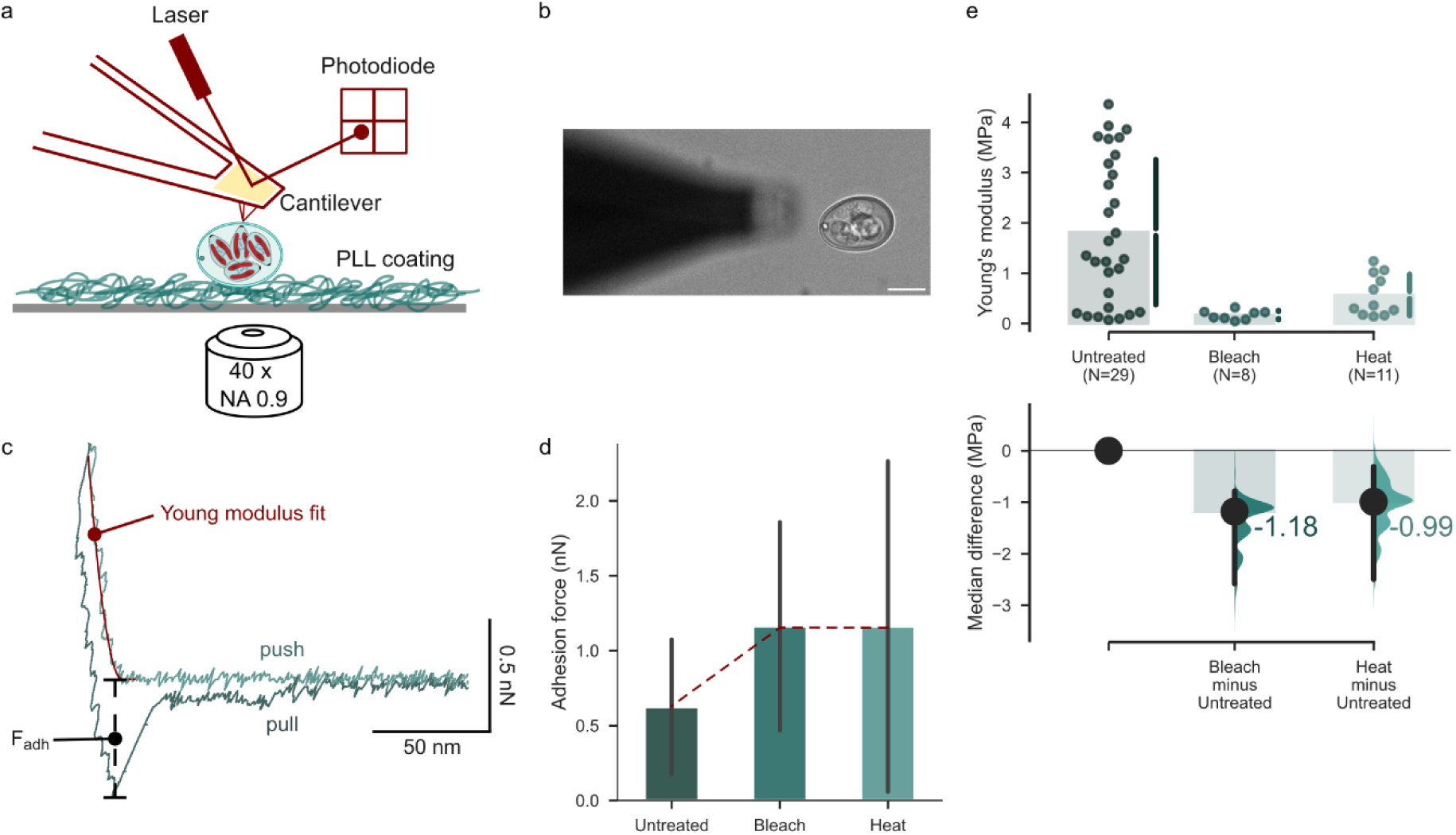
| Atomic force microscopy and microindentation reveal treatment-dependent changes in oocyst wall mechanics. **a**, Schematic representation of the atomic force microscopy (AFM) setup used to probe oocyst wall mechanics. Oocysts were immobilized on a poly-L-lysine (PLL)-coated surface and indented with an AFM cantilever; cantilever deflection was detected by a laser–photodiode system. **b**, Representative bright-field micrograph of an oocyst positioned near the AFM cantilever. Scale bar, 10 µm. **c**, Representative AFM force–distance curve showing cantilever approach and retraction. The Young’s modulus was extracted from the indentation fit during the approach phase, and the adhesion force was measured during cantilever retraction. **d**, Adhesion force measured for untreated, bleach-treated and heat-treated oocysts. Bars indicate the mean and error bars indicate s.d. **e**, Young’s modulus of untreated, bleach-treated and heat-treated oocysts. Each dot represents an individual measurement; bars indicate the group summary. The lower estimation plot shows the median difference between treated and untreated oocysts.

**Supplementary Fig. 4.**
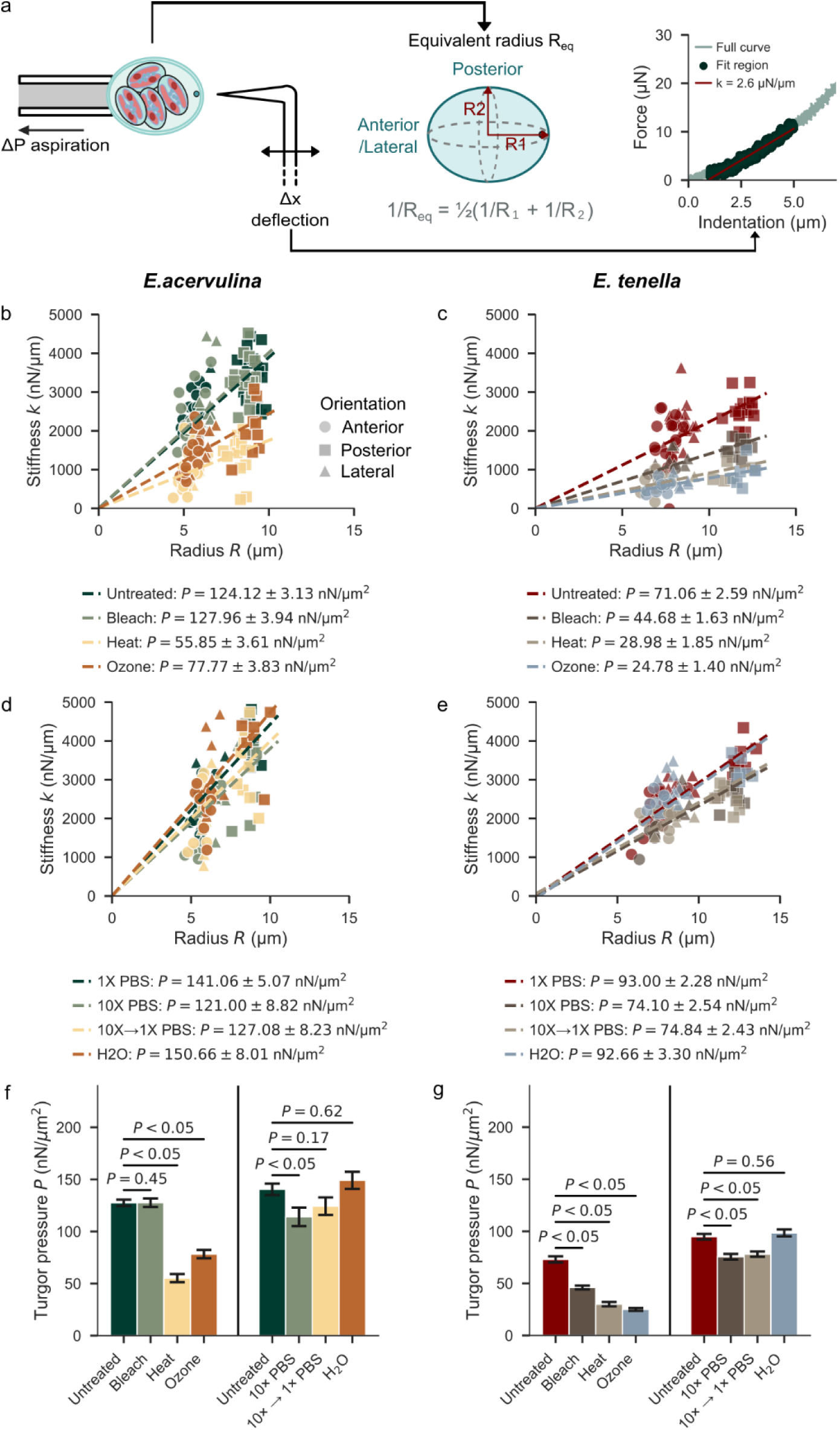
| Radius-dependent stiffness reveals treatment-induced changes in oocyst internal pressure. **a**, Schematic of the microindentation-based approach used to estimate oocyst internal pressure. Oocyst geometry was described by an equivalent radius, *Req*, calculated from the anterior/posterior and lateral radii. The contact stiffness, *k*, was extracted from the linear fit of the force–indentation curve. **b,c**, Relationship between contact stiffness, *k*, and equivalent radius, *Req*, in *E. acervulina* (**b**) and *E. tenella* (**c**) oocysts after untreated, bleach, heat and ozone treatments. **d,e**, Relationship between stiffness, *k*, and equivalent radius, *Req*, in *E. acervulina* (**d**) and *E. tenella* (**e**) oocysts under osmotic conditions: 1× PBS, 10× PBS, 10× PBS → 1× PBS and H₂O. Marker shape indicates indentation orientation. Dashed lines indicate linear fits used to estimate internal pressure. **f,g**, Estimated internal pressure in *E. acervulina* (**f**) and *E. tenella* (**g**) oocysts across disinfectant and osmotic treatments. Bars indicate pressure values derived from the stiffness–radius fits; error bars indicate the uncertainty of the fitted estimate. *P* values were calculated using two-sided Mann–Whitney *U* tests for the indicated pairwise comparisons.

**Supplementary Table 1.** | Effect sizes for all pairwise comparisons against untreated controls. Each treatment was compared with the corresponding untreated control in *E. acervulina* and *E. tenella* for oocyst area, wall autofluorescence (AF), and rupture force at lateral/posterior (L/P) and anterior (A) indentation positions. Columns give the median and number of oocysts (n) for untreated and treated conditions, the two-tailed Mann–Whitney *U*-test *P* value, Cliff’s δ (non-parametric effect size), and the effect-size magnitude. Magnitudes: negligible (|δ| < 0.147), small (0.147 ≤ |δ| < 0.33), medium (0.33 ≤ |δ| < 0.474), large (|δ| ≥ 0.474). Treatments are grouped by assay (digestive factors, disinfectants, osmotic conditions). δ ranges from −1 to +1; a positive value indicates an increase relative to untreated and a negative value a decrease, with magnitude reflecting the degree of distributional non-overlap.

| *Species* | *Assay* | *Measure* | *Treatment* | *Untreated, median (n)* | *Condition, median (n)* | *P value* | *Cliff's $\Delta$* | *Effect magnitude* |
| --- | --- | --- | --- | --- | --- | --- | --- | --- |
| *E. acervulina* | Digestive | Area | Trypsin | 196.7 (177) | 197.2 (129) | 0.58 | 0.04 | Negligible |
| *E. acervulina* | Digestive | Area | Bile salts | 196.7 (177) | 205.3 (88) | 0.02 | 0.18 | Small |
| *E. acervulina* | Digestive | Area | Trypsin+bile salts | 196.7 (177) | 213.4 (41) | 0.01 | 0.28 | Small |
| *E. tenella* | Digestive | Area | Trypsin | 301.8 (122) | 299.8 (30) | 0.51 | -0.08 | Negligible |
| *E. tenella* | Digestive | Area | Bile salts | 301.8 (122) | 311 (64) | 0.21 | 0.11 | Negligible |
| *E. tenella* | Digestive | Area | Trypsin+bile salts | 301.8 (122) | 308.3 (106) | 0.39 | 0.07 | Negligible |
| *E. acervulina* | Disinfectant | Area | Bleach | 200.1 (466) | 187.7 (170) | <0.01 | -0.21 | Small |
| *E. acervulina* | Disinfectant | Area | Heat | 200.1 (466) | 177.6 (111) | <0.01 | -0.45 | Medium |
| *E. acervulina* | Disinfectant | Area | Ozone | 200.1 (466) | 198.2 (50) | 0.58 | -0.05 | Negligible |
| *E. tenella* | Disinfectant | Area | Bleach | 347.2 (159) | 344.6 (74) | 0.89 | -0.01 | Negligible |
| *E. tenella* | Disinfectant | Area | Heat | 347.2 (159) | 314.9 (93) | <0.01 | -0.4 | Medium |
| *E. tenella* | Disinfectant | Area | Ozone | 347.2 (159) | 303.8 (55) | <0.01 | -0.44 | Medium |
| *E. acervulina* | Osmolarity | Area | PBS10X | 198.7 (177) | 194.6 (59) | 0.85 | 0.02 | Negligible |
| *E. acervulina* | Osmolarity | Area | 10×→1×PBS | 198.7 (177) | 198.2 (45) | 0.66 | 0.04 | Negligible |
| *E. acervulina* | Osmolarity | Area | H2O | 198.7 (177) | 196 (58) | 0.94 | -0.01 | Negligible |
| *E. tenella* | Osmolarity | Area | PBS10X | 301.8 (122) | 312.6 (60) | 0.34 | 0.09 | Negligible |
| *E. tenella* | Osmolarity | Area | 10×→1×PBS | 301.8 (122) | 312.6 (53) | 0.09 | 0.16 | Small |
| *E. tenella* | Osmolarity | Area | H2O | 301.8 (122) | 308.1 (94) | 0.47 | 0.06 | Negligible |
| *E. acervulina* | Digestive | AF | Trypsin | 1672.9 (177) | 1442.8 (129) | <0.01 | -0.32 | Small |
| *E. acervulina* | Digestive | AF | Bile salts | 1672.9 (177) | 1509.2 (88) | 0.09 | -0.13 | Negligible |
| *E. acervulina* | Digestive | AF | Trypsin+bile salts | 1672.9 (177) | 1237.3 (41) | <0.01 | -0.59 | Large |
| *E. tenella* | Digestive | AF | Trypsin | 2426.7 (122) | 2522.7 (30) | 0.15 | 0.17 | Small |
| *E. tenella* | Digestive | AF | Bile salts | 2426.7 (122) | 2458 (64) | 0.9 | 0.01 | Negligible |
| *E. tenella* | Digestive | AF | Trypsin+bile salts | 2426.7 (122) | 2590.3 (106) | 0.01 | 0.2 | Small |
| *E. acervulina* | Disinfectant | AF | Bleach | 1534 (229) | 525.2 (35) | <0.01 | -0.93 | Large |
| *E. acervulina* | Disinfectant | AF | Heat | 1534 (229) | 1155.6 (123) | <0.01 | -0.53 | Large |
| *E. acervulina* | Disinfectant | AF | Ozone | 1534 (229) | 414.8 (48) | <0.01 | -0.98 | Large |
| *E. tenella* | Disinfectant | AF | Bleach | 1871.1 (158) | 1331.7 (74) | <0.01 | -0.58 | Large |
| *E. tenella* | Disinfectant | AF | Heat | 1871.1 (158) | 2886.8 (93) | <0.01 | 0.84 | Large |
| *E. tenella* | Disinfectant | AF | Ozone | 1871.1<br>(158) | 567.4 (55) | <0.01 | -0.98 | Large |
| *E. acervulina* | Osmolarity | AF | PBS10X | 1672.9<br>(177) | 1454.7 (59) | <0.01 | -0.36 | Medium |
| *E. acervulina* | Osmolarity | AF | 10×→1×PBS | 1672.9<br>(177) | 1339.4 (45) | <0.01 | -0.48 | Large |
| *E. acervulina* | Osmolarity | AF | H2O | 1672.9<br>(177) | 1453.6 (58) | <0.01 | -0.3 | Small |
| *E. tenella* | Osmolarity | AF | PBS10X | 2426.7<br>(122) | 2310.4 (60) | 0.97 | 0 | Negligible |
| *E. tenella* | Osmolarity | AF | 10×→1×PBS | 2426.7<br>(122) | 2583 (53) | <0.01 | 0.27 | Small |
| *E. tenella* | Osmolarity | AF | H2O | 2426.7<br>(122) | 2781.3 (94) | 0 | 0.35 | Medium |
| *E. acervulina* | Digestive | Force<br>(L/P) | Trypsin | 34.8 (25) | 31.7 (20) | 0.23 | -0.21 | Small |
| *E. acervulina* | Digestive | Force<br>(L/P) | Bile salts | 34.8 (25) | 31.5 (20) | 0.34 | -0.17 | Small |
| *E. acervulina* | Digestive | Force<br>(L/P) | Trypsin+bile<br>salts | 34.8 (25) | 26.8 (20) | 0.01 | -0.45 | Medium |
| *E. acervulina* | Digestive | Force (A) | Trypsin | 25.3 (11) | 18.9 (10) | 0.22 | -0.33 | Small |
| *E. acervulina* | Digestive | Force (A) | Bile salts | 25.3 (11) | 15.4 (10) | 0.02 | -0.64 | Large |
| *E. acervulina* | Digestive | Force (A) | Trypsin+bile<br>salts | 25.3 (11) | 19.7 (10) | 0.05 | -0.51 | Large |
| *E. tenella* | Digestive | Force<br>(L/P) | Trypsin | 21.2 (22) | 19.5 (20) | 0.59 | -0.1 | Negligible |
| *E. tenella* | Digestive | Force<br>(L/P) | Bile salts | 21.2 (22) | 22.6 (20) | 0.18 | 0.25 | Small |
| *E. tenella* | Digestive | Force<br>(L/P) | Trypsin+bile<br>salts | 21.2 (22) | 24 (19) | 0.01 | 0.48 | Large |
| *E. tenella* | Digestive | Force (A) | Trypsin | 15.9 (11) | 9.4 (10) | 0.07 | -0.47 | Medium |
| *E. tenella* | Digestive | Force (A) | Bile salts | 15.9 (11) | 3.8 (10) | 0.02 | -0.64 | Large |
| *E. tenella* | Digestive | Force (A) | Trypsin+bile<br>salts | 15.9 (11) | 8.8 (9) | 0.07 | -0.49 | Large |
| *E. acervulina* | Disinfectant | Force<br>(L/P) | Bleach | 32.8 (29) | 32.6 (36) | 0.49 | -0.1 | Negligible |
| *E. acervulina* | Disinfectant | Force<br>(L/P) | Heat | 32.8 (29) | 22.7 (30) | <0.01 | -0.66 | Large |
| *E. acervulina* | Disinfectant | Force (L/P) | Ozone | 32.8 (29) | 24.8 (27) | <0.01 | -0.62 | Large |
| *E. acervulina* | Disinfectant | Force (A) | Bleach | 24.8 (25) | 26.1 (27) | 0.74 | -0.05 | Negligible |
| *E. acervulina* | Disinfectant | Force (A) | Heat | 24.8 (25) | 6.3 (21) | <0.01 | -0.97 | Large |
| *E. acervulina* | Disinfectant | Force (A) | Ozone | 24.8 (25) | 22.4 (15) | 0.05 | -0.37 | Medium |
| *E. tenella* | Disinfectant | Force (L/P) | Bleach | 21.2 (17) | 20.2 (22) | 0.06 | -0.36 | Medium |
| *E. tenella* | Disinfectant | Force (L/P) | Heat | 21.2 (17) | 23.3 (14) | 0.56 | 0.13 | Negligible |
| *E. tenella* | Disinfectant | Force (L/P) | Ozone | 21.2 (17) | 21.2 (20) | 0.59 | -0.11 | Negligible |
| *E. tenella* | Disinfectant | Force (A) | Bleach | 17 (10) | 15.8 (5) | 0.68 | -0.16 | Small |
| *E. tenella* | Disinfectant | Force (A) | Heat | 17 (10) | 5.9 (10) | 0.19 | -0.36 | Medium |
| *E. tenella* | Disinfectant | Force (A) | Ozone | 17 (10) | 15 (10) | 0.43 | -0.22 | Small |
| *E. acervulina* | Osmolarity | Force (L/P) | PBS10X | 29.5 (39) | 37.6 (20) | 0.14 | 0.24 | Small |
| *E. acervulina* | Osmolarity | Force (L/P) | 10×→1×PBS | 29.5 (39) | 33.4 (20) | 0.63 | 0.08 | Negligible |
| *E. acervulina* | Osmolarity | Force (L/P) | H2O | 29.5 (39) | 28.8 (23) | 0.14 | -0.23 | Small |
| *E. acervulina* | Osmolarity | Force (A) | PBS10X | 23.5 (20) | 27.6 (10) | 0.95 | -0.02 | Negligible |
| *E. acervulina* | Osmolarity | Force (A) | 10×→1×PBS | 23.5 (20) | 25.8 (10) | 0.5 | 0.16 | Small |
| *E. acervulina* | Osmolarity | Force (A) | H2O | 23.5 (20) | 20.4 (11) | 0.17 | -0.31 | Small |
| *E. tenella* | Osmolarity | Force (L/P) | PBS10X | 21.2 (22) | 24.6 (21) | <0.01 | 0.57 | Large |
| *E. tenella* | Osmolarity | Force (L/P) | 10×→1×PBS | 21.2 (22) | 25.5 (20) | <0.01 | 0.65 | Large |
| *E. tenella* | Osmolarity | Force (L/P) | H2O | 21.2 (22) | 22.6 (20) | 0.05 | 0.36 | Medium |
| *E. tenella* | Osmolarity | Force (A) | PBS10X | 15.9 (11) | 21.2 (10) | 0.22 | 0.33 | Small |
| *E. tenella* | Osmolarity | Force (A) | 10×→1×PBS | 15.9 (11) | 18.8 (10) | 0.13 | 0.4 | Medium |
| *E. tenella* | Osmolarity | Force (A) | H2O | 15.9 (11) | 20 (10) | 0.11 | 0.42 | Medium |

## References

1. Tenter, A. M. et al. The conceptual basis for a new classification of the coccidia. Int J Parasitol 32, 595–616 (2002).

2. Berto, B. P., McIntosh, D. & Lopes, C. W. G. Studies on coccidian oocysts (Apicomplexa: Eucoccidiorida). Rev Bras Parasitol Vet 23, 1–15 (2014).

3. Belli, S. I., Smith, N. C. & Ferguson, D. J. P. The coccidian oocyst: a tough nut to crack! Trends Parasitol 22, 416–423 (2006).

4. McCaughan, K. J. & Kniel, K. E. Current Knowledge and Future Directions for Cyclospora cayetanensis Research and Its Surrogates. Compr Rev Food Sci Food Saf 25, e70327 (2026).

5. Robert-Gangneux, F. & Dardé, M.-L. Epidemiology of and diagnostic strategies for toxoplasmosis. Clin Microbiol Rev 25, 264–296 (2012).

6. Blake, D. P., Marugan-Hernandez, V. & Tomley, F. M. Spotlight on avian pathology: Eimeria and the disease coccidiosis. Avian Pathol 1–5 (2021).

7. Innes, E. A., Hamilton, C., Garcia, J. L., Chryssafidis, A. & Smith, D. A one health approach to vaccines against Toxoplasma gondii. Food Waterborne Parasitol 15, e00053 (2019).

8. Shapiro, K. et al. Environmental transmission of Toxoplasma gondii: Oocysts in water, soil and food. Food Waterborne Parasitol 15, e00049 (2019).

9. Daugschies, A., Bangoura, B. & Lendner, M. Inactivation of exogenous endoparasite stages by chemical disinfectants: current state and perspectives. Parasitol Res 112, 917–932 (2013).

10. Dumètre, A. & Dardé, M. L. How to detect Toxoplasma gondii oocysts in environmental samples? FEMS Microbiol Rev 27, 651–661 (2003).

11. Jenkins, M. C., Parker, C., O’Brien, C., Miska, K. & Fetterer, R. Differing susceptibilities of Eimeria acervulina, Eimeria maxima, and Eimeria tenella oocysts to desiccation. J Parasitol 99, 899–902 (2013).

12. Freppel, W. et al. Structure, composition, and roles of the Toxoplasma gondii oocyst and sporocyst walls. Cell Surf 5, 100016 (2019).

13. Bushkin, G. G. et al. Evidence for a structural role for acid-fast lipids in oocyst walls of Cryptosporidium, Toxoplasma, and Eimeria. mBio 4, e00387–00313 (2013).

14. Mai, K. et al. Oocyst wall formation and composition in coccidian parasites. Mem Inst Oswaldo Cruz 104, 281–289 (2009).

15. Samuelson, J., Bushkin, G. G., Chatterjee, A. & Robbins, P. W. Strategies To Discover the Structural Components of Cyst and Oocyst Walls. Eukaryot Cell 12, 1578–1587 (2013).

16. Belli, S. I., Wallach, M. G., Luxford, C., Davies, M. J. & Smith, N. C. Roles of Tyrosine-Rich Precursor Glycoproteins and Dityrosine- and 3,4-Dihydroxyphenylalanine-Mediated Protein Cross-Linking in Development of the Oocyst Wall in the Coccidian Parasite Eimeria maxima. Eukaryot Cell 2, 456–464 (2003).

17. Dumètre, A. et al. Mechanics of the Toxoplasma gondii oocyst wall. Proc Natl Acad Sci U S A 110, 11535–11540 (2013).

18. Bushkin, G. G. et al. β-1,3-glucan, which can be targeted by drugs, forms a trabecular scaffold in the oocyst walls of Toxoplasma and Eimeria. mBio 3, e00258–12 (2012).

19. Municio-Diaz, C. et al. Mechanobiology of the cell wall – insights from tip-growing plant and fungal cells. J Cell Sci 135, jcs259208 (2022).

20. Popov, V. L. Viscoelastic Properties of Elastomers. in Contact Mechanics and Friction: Physical Principles and Applications (ed. Popov, V. L.) 231–253 (Springer, 2010).

21. Augendre, L., et al. Surrogates of foodborne and waterborne protozoan parasites: A review. Food Waterborne Parasitol 33, e00212 (2023).

22. Burrell, A., Tomley, F. M., Vaughan, S. & Marugan-Hernandez, V. Life cycle stages, specific organelles and invasion mechanisms of Eimeria species. Parasitology 147, 263–278 (2019).

23. Rose, M. E. & Hesketh, P. Eimeria tenella: localization of the sporozoites in the caecum of the domestic fowl. Parasitology 102, 317–324 (1991).

24. Augendre, L., et al. *Eimeria acervulina* is a promising surrogate for *Toxoplasma gondii* oocysts exposed to chemical and physical treatments. Exp Parasitol 278, 109049 (2025).

25. Dumètre, A., Dubey, J. P. & Ferguson, D. J. P. Effect of household bleach on the structure of the sporocyst wall of Toxoplasma gondii. Parasite 28, 68 (2021).

26. Fuller, A. & McDougald, L. Lectin-binding by sporozoites of Eimeria tenella. Parasitol Res 88, 118–125 (2002).

27. Michalski, W. P., Prowse, S. J., Bacic, A. & Fahey, K. J. Molecular characterisation of peanut agglutinin-binding glycoproteins from *Eimeria tenella*. Int J Parasitol 23, 985–995 (1993).

28. El Husseiny, J., Puech, P.-H., Dumètre, A. & Husson, J. Protocol for quantifying coccidian parasite mechanics and rupture force using advanced micromanipulation-based piercing. STAR Protoc 6, 104234 (2025).

29. Vella, D., Ajdari, A., Vaziri, A. & Boudaoud, A. The indentation of pressurized elastic shells: from polymeric capsules to yeast cells. J R Soc Interface 9, 448–455 (2012).

30. Doran, D. J. & Farr, M. M. Excystation of the Poultry Coccidium, Eimeria acervulina. J Protozool 9, 154–161 (1962).

31. Cha, J.-O., Talha, A. F. S. M., Lim, C. W. & Kim, B. Effects of glass bead size, vortexing speed and duration on Eimeria acervulina oocyst excystation. Exp Parasitol 138, 18–24 (2014).

32. Wiedmer, S. et al. New insights into the excystation process and oocyst morphology of rodent Eimeria species. Protist 162, 668–678 (2011).

33. Relat-Goberna, J., Beedle, A. E. M. & Garcia-Manyes, S. The Nanomechanics of Lipid Multibilayer Stacks Exhibits Complex Dynamics. Small 13, 1700147 (2017).

34. Sen, S., Subramanian, S. & Discher, D. E. Indentation and Adhesive Probing of a Cell Membrane with AFM: Theoretical Model and Experiments. Biophys J 89, 3203–3213 (2005).

35. Mashmoushy, H., Zhang, Z. & Thomas, C. R. Micromanipulation measurement of the mechanical properties of baker’s yeast cells. Biotechnol Tech 12, 925–929 (1998).

36. Haas, S., Körner, S., Zintel, L. & Hubbuch, J. Changing mechanical properties of photopolymerized, dityrosine-crosslinked protein-based hydrogels. Front Bioeng Biotechnol 10, 1006438 (2022).

37. Zheng, D., Lin, S., Ni, J. & Zhao, X. Fracture and fatigue of entangled and unentangled polymer networks. Extreme Mech Lett 51, 101608 (2022).

38. Gérard, C. et al. Inactivation of parasite transmission stages: Efficacy of treatments on foods of non-animal origin. TIFS 91, 12–23 (2019).

39. Cha, J. O. et al. Oocyst-Shedding Patterns of Three Eimeria Species in Chickens and Shedding Pattern Variation Depending on the Storage Period of Eimeria tenella Oocysts. J Parasitol 104, 18–22 (2018).

40. Répérant, J.-M., Thomas-Hénaff, M., Benoit, C., Le Bihannic, P. & Eterradossi, N. The impact of maturity on the ability of Eimeria acervulina and Eimeria meleagrimitis oocysts to sporulate. Parasite 28, 32 (2021).

41. Ribeiro E Silva, A., et al. Genome-Wide Expression Patterns of Rhoptry Kinases during the Eimeria tenella Life-Cycle. Microorganisms 9, 1621 (2021).

42. Edelstein, A. D. et al. Advanced methods of microscope control using μManager software. J Biol Methods 1, e10 (2014).

43. Schmidt, U., Weigert, M., Broaddus, C. & Myers, G. Cell Detection with Star-Convex Polygons. in Medical Image Computing and Computer Assisted Intervention – MICCAI 2018 (eds Frangi, A. F., Schnabel, J. A., Davatzikos, C., Alberola-López, C. & Fichtinger, G.) 265–273 (Springer International Publishing, 2018).

44. Weigert, M. et al. Content-aware image restoration: pushing the limits of fluorescence microscopy. Nat Methods 15, 1090–1097 (2018).

45. Arzt, M., et al. LABKIT: Labeling and Segmentation Toolkit for Big Image Data. Front Comput Sci 4, (2022).

46. Torro, R. et al. Celldetective: an AI-enhanced image analysis tool for unraveling dynamic cell interactions. eLife 14, (2025).

47. Husson, J. Measuring Cell Mechanical Properties Using Microindentation. in Mechanobiology: Methods and Protocols (ed. Zaidel-Bar, R.) 3–23 (Springer US, 2023).

48. Johnson, J. & Reid, W. M. Anticoccidial drugs: Lesion scoring techniques in battery and floor-pen experiments with chickens. Exp Parasitol 28, 30–36 (1970).

49. Cliff, N. Dominance statistics: Ordinal analyses to answer ordinal questions. Psychological Bulletin 114, 494–509 (1993).

50. Romano, J., Kromrey, J. D., Coraggio, J. & Skowronek, J. Appropriate statistics for ordinal level data: Should we really be using t-test and Cohen’s d for evaluating group differences on the NSSE and other surveys. in annual meeting of the Florida Association of Institutional Research vol. 177 (2006).

